# Continuum architecture dynamics of vesicle tethering in exocytosis

**DOI:** 10.1101/2025.02.05.635468

**Authors:** Marta Puig-Tintó, Sebastian Ortiz, Sasha Meek, Raffaele Coray, Altair C. Hernández, Anna Castellet, Eric Kramer, Laura I. Betancur, Philipp Hoess, Markus Mund, Mercè Izquierdo-Serra, Baldo Oliva, Alex de Marco, Jonas Ries, Daniel Castaño-Díez, Carlo Manzo, Oriol Gallego

## Abstract

Essential for eukaryotic organisms, multiple copies of the exocyst complex tether each secretory vesicle to the plasma membrane (PM) in constitutive exocytosis. The exocyst higher-order structure (ExHOS) that coordinates the action of these multiple exocysts remains unexplored. We integrated particle-tracking, super-resolution microscopy and cryo-electron tomography in a model that time-resolves the continuum conformational landscape of the ExHOS and functionally annotates its different conformations. We found that 7 exocysts form flexible ring-shaped ExHOS that tether vesicles at <45 nm from the PM. The ExHOS, initially 19 nm in radius, rapidly expands while it pulls the vesicle towards the PM. Subsequently, the ExHOS stabilizes, securing the vesicle at ∼4 nm from the PM. After fusion, Sec18 mediates the ExHOS disassembly when its radius is 38 nm. By resolving the fundamental biophysical principles of tethering we bridged the gap between static isolated structures and the dynamic and multimeric nature of exocytosis.

## INTRODUCTION

Constitutive exocytosis (hereafter exocytosis), the uninterrupted transport of secretory vesicles to and subsequent fusion with the plasma membrane (PM), is an essential cellular process for nearly all eukaryotes. Exocytosis is critical for the preservation of PM homeostasis, cell growth and cell division. At the heart of this intricate process lies the tethering, a finely tuned event that is responsible for the specific docking of cargo-loaded vesicles with the target PM. The exocyst, a conserved heterooctameric protein complex, is the main component of tethering^1–3^. Exocyst function relies on the interplay with other proteins that actively regulate exocytosis. For example, exocyst function requires the interaction, at the vesicle surface, with the nucleotide guanosine triphosphatase Sec4 and its guanyl-nucleotide exchange factor Sec2^4,5^. The exocyst also binds Sec9, a soluble N-ethylmaleimide-sensitive factor attachment protein receptor (SNARE). Exocyst-Sec9 interaction supports the exocytic *trans*-SNARE complex formation, which is critical for subsequent vesicle fusion^6^. After cargo release, the AAA ATPase Sec18 hydrolyses ATP and, together with the α-SNAP co-chaperone Sec17, it culminates the process of exocytosis by disassembling the SNARE complex, which is recycled for the progression of a new round of tethering and fusion^7,8^.

*In vitro* experiments have been essential to provide high-resolution data of this tethering complex. The most complete structure of the isolated complex measures 32 nm long and 13 nm wide. Although this structure still lacks 18% of the complex, it uncovered the atomic details that sustain the assembly of the octamer in two tetrameric modules: Module I contains Sec3, Sec5, Sec6 and Sec8 subunits, while Module II consists of Sec10, Sec15, Exo70 and Exo84 subunits^9,10^. *In vitro* experiments also located the regions in Sec15 and Sec6 C-termini that bind functional partners such as Sec4-Sec2 and Sec9, respectively^10,11^. However, in the cell, the tethering of a single vesicle involves multiple copies of the exocyst^12–14^ with defined stoichiometries and temporal correlation with its partners^12,14^. Therefore, tethering is governed by a dynamic higher-order mechanism, which could not be reconstituted *in vitro*. The analysis *in situ* of the higher-order mechanism of tethering is further complicated by the polarized and transient nature of the exocytic events, which tend to occur in highly localized regions of the PM^14^. These challenges, combined with the intrinsic complexity and constitutive dynamism of exocytosis, have limited the quantitative studies of exocytosis under native conditions. Thus, the spatio-temporal organization of the multiple exocysts mediating vesicle tethering (the exocyst higher-order structure, hereafter named ExHOS), along with its functional annotation, remains as an unresolved milestone in the field. Using *Saccharomyces cerevisiae*, we integrated 2-color simultaneous particle tracking of exocytic proteins, super-resolution microscopy and cryo-correlative light and electron microscopy (cryo-CLEM) to resolve the dynamic architecture of multiple exocysts and the secretory vesicle during exocytosis.

## RESULTS

### Measuring the vesicle radius and the exocyst copy number to model the tethering

Exocytosis inherently involves the interaction and shaping of membranes (secretory vesicles and PM). The combination of focused ion beam scanning electron microscopy (FIB-SEM) and cryo-electron tomography (cryo-ET) offers an intimate 3D view of this membrane ultrastructure in vitrified cells. We imaged and segmented 175 vesicles adjacent (i.e., <70 nm of distance) to the PM of yeast cells (Figures 1A and S1). Spherical vesicles, with an average radius of 41 ± 4 nm, locate preferentially at distances below 45 nm from the PM. This suggests the presence of an underlying mechanism that drives this proximal accumulation, such as exocyst-mediated tethering (Figure 1A).

**Figure 1.**
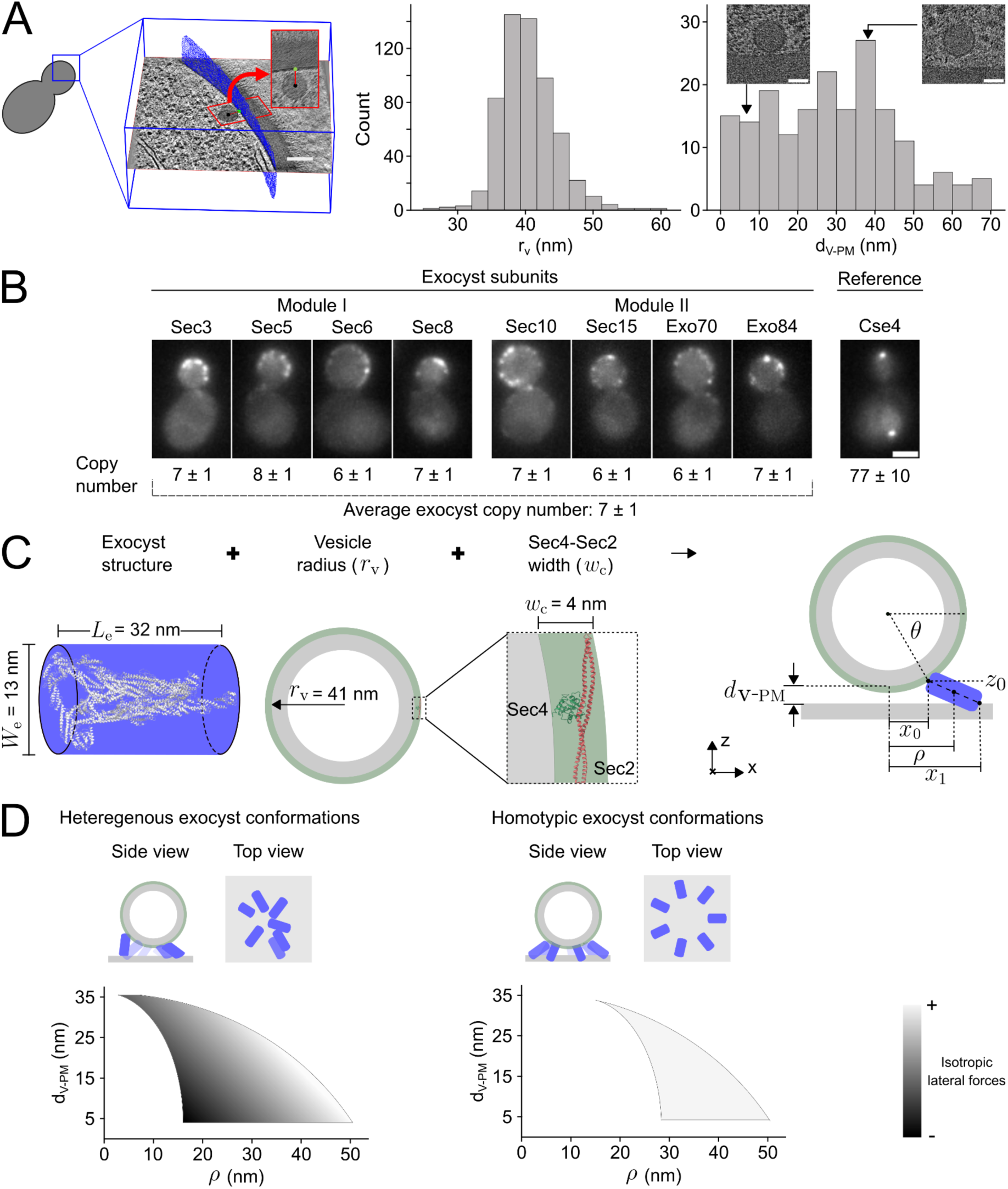
Modeling the spatial arrangement of vesicle tethering. (A) Yeast PM was imaged by cryo-ET. Upon segmentation of secretory vesicles and the PM (left and Figure S1; scale bar: 100 nm), the vesicle radius (r_v_, center) and the shortest distance between the vesicle and the PM (d_V-PM_, right) were measured. Illustrative tomogram slices are shown (scale bars: 50 nm). (B) Average copy number of each subunit in exocyst clusters was measured using Cse4-GFP as reference (Figure S2). Illustrative images of cells expressing the indicated subunit fused to GFP. Data is shown as mean ± pooled SEM. Scale bar: 2 μm. (C) Illustration of the constraints used to model possible ExHOS (Figure S3 and Methods). The exocyst is modeled as a cylinder fitting the dimensions of the cryo-EM structure (left)^10^. The vesicle is modeled as a 41 nm radius sphere (center) with additional width accounting for the Sec4-Sec2 complex (PDB: 2OCY). (D) The possible conformational space of the ExHOS was modeled considering two scenarios: when the 6-8 exocysts in a cluster adopt heterogeneous conformations (left) and when the 6-8 exocysts arrange in a homotypic conformation (right). The gray scale illustrates the suitability of modeled ExHOS to apply isotropic lateral forces.

The first step in elucidating the higher-order mechanism of such vesicle tethering was to determine the stoichiometry of exocysts involved in this process, which remains inconclusive and incomplete for all exocyst subunits^12–14^. We measured the average number of each subunit fused to GFP using a live-cell fluorescence intensity ratiometric assay^15^, with Cse4-GFP as a reference (Methods)^16^. In agreement with the number of Sec3, Sec5, Sec6, Sec8 and Exo70 subunits found in exocyst clusters of mammalian cells^12^, we found that 7 ± 1 (mean ± pooled SEM) exocysts are present at exocytic events, on average, with the expected 1 to 1 stoichiometry for all the subunits (Figures 1B and S2).

Subsequently, we investigated how 6 to 8 exocysts can tether vesicles situated at different distances from the PM. To address this question, we modeled the possible conformational space of the ExHOS that can bind the vesicle and the PM simultaneously (Figure 1C and Methods). Each of the exocyst was represented by a cylinder of 13 nm of radius and 32 nm of height. The radial location of a given exocyst (*ρ*), which is the distance of the exocyst centroid to the central axis in the vesicle-PM interface, was used henceforth as a descriptor of the ExHOS (Figure 1C, right). Given the fixed length of the exocyst in our model, *ρ* depends on the vesicle-PM distance (d_V-PM_ in Figure 1C) and the conformation that the exocyst adopts in relation to the PM plane.

The conformational relationship among the multiple exocysts involved in the ExHOS imposes two plausible architectural modalities linked to different mechanisms of tethering. A first mechanism, where the multiple exocysts tethering a vesicle adopt heterogenous conformations, would allow the 6 to 8 exocysts to be unevenly dispersed along a patch-shaped ExHOS between the vesicle and the PM. Hence, exocysts positions could range from a *ρ* of nearly 6.5 nm, corresponding to the cylinder radius, to 50 nm (Figure 1D, left). A second mechanism is also plausible where the exocysts conformation is synchronized. By requiring that all the exocysts adopt the same conformation relative to the vesicle and the PM, ring-shaped models are obtained, where all exocysts share the same *ρ* value (hereafter, ExHOS radius) (Figure 1D, right). The radius of the ring-shaped ExHOS depends on the number of complexes that must be accommodated and the distance at which the vesicle is tethered to the PM (Figure S3). Thus, the theoretical ExHOS with 6 exocysts tethering the vesicle at 34 nm from the PM has a radius of 15 nm, while the ExHOS tethering the vesicle at 4 nm from the PM has a radius of up to 50 nm (Figure 1D, right). The biological relevance of both mechanisms might be related with the distribution of the forces exerted by the exocysts to control the lateral diffusion of the vesicle. Thus, ring-shaped ExHOS are more likely to provide isotropic lateral forces that might result in a vesicle displacement orthogonal to the PM.

### Exocysts organize in different classes of higher-order structures

To experimentally assess the ExHOS *in situ* we used SMLM, a technique that provides ∼20 nm resolution. We imaged fixed cells endogenously expressing exocyst subunits Sec5, Sec6 and Sec8, from Module I, fused to the photoswitchable fluorescent protein mMaple (mM), and Exo84, from Module II, fused to GFP (exocyst-mM-GFP). We obtained two-dimensional projections of exocyst-mM-GFP clusters by setting the focal plane at the bottom of the cells (Figure 2A). Correlating SMLM and diffraction-limited wide-field microscopy (DL) images allowed us to segment 136 individual exocyst-mM-GFP clusters (Figure 2A). Due to its sensitivity and resolution, SMLM has been demonstrated to be a robust approach to quantify the average number of protein clusters^16^. Consistent with the live-cell fluorescence intensity ratiometric assay, on average, 5.8 ± 3.4 exocysts populate exocyst-mM-GFP clusters (Figure 2A and S4).

**Figure 2.**
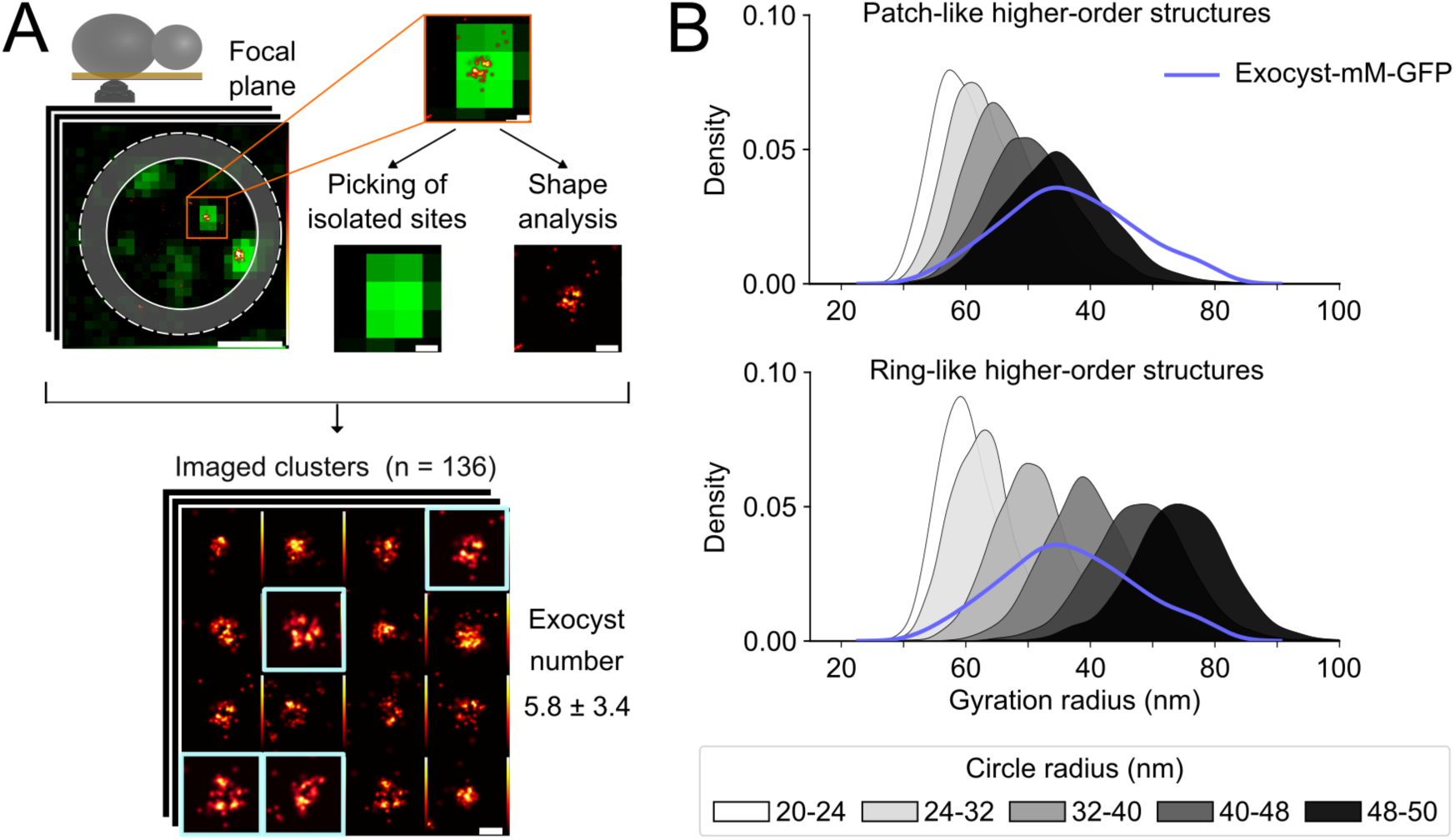
Structural analysis of the ExHOS by SMLM. (A) Yeast cells expressing exocyst-mM-GFP were imaged at the bottom of the cell. SMLM and diffraction-limited (DL) images were correlated. A representative SMLM-DL image of a single cell is shown (top). The DL signal was used to identify 136 isolated exocyst clusters. A mosaic of representative images is shown with ring-shaped ExHOS highlighted in white squares (bottom). The exocyst copy number (mean ± pooled SD) was estimated using Nup188-mM as reference (Figure S4 and Methods). Errors were calculated from three biological replicates. Scale bars: 100 nm. (B) Circle-shaped ExHOSs of varying radius (gray scale) and populated by 6 to 8 exocysts were considered, with two possible scenarios: when exocysts are distributed inside the entire circle (patch-like, top) and when the exocysts are located along the periphery of a circle (ring-like, bottom). Only the ring-shaped ExHOS can explain the distribution of gyration radius of localizations observed in the experimental data (blue line).

The sparsity of exocysts populating ExHOS makes its structural analysis by SMLM highly sensitive to the uneven photoconversion of mMaple molecules and its limited labeling efficiency^17^. Nonetheless, in agreement with a homotypic exocyst conformation in ExHOS, visual inspection identified ring-like shapes in some of the exocyst-mM-GFP cluster images (Figure 2A). To tackle the conundrum of the ExHOS shape more systematically, we compiled a synthetic dataset of the expected SMLM images resulting from imaging exocyst-mM clusters where the multiple exocyst follow heterogeneous (i.e., patch-like) and homotypic (i.e., ring-like) ExHOS. We considered radii between 20 and 50 nm (Methods). We then compared the gyration radius, a measure of dispersion, for the localizations observed in the experimental dataset, with the gyration radius of the simulated data. Interestingly, only the synthetic dataset of ring-shaped ExHOS could recapitulate the localizations’ gyration radius above 70 nm observed experimentally (Figure 2B).

In addition, the simulations also indicate that the coexistence of ring-shaped ExHOS with different radii is necessary to fully explain the broad range of localizations’ gyration radius observed experimentally.

Cryo-ET data, computational predictions and SMLM-DL imaging imply that 7 exocysts on average can arrange in a ring-shaped ExHOS that adopts different radii on the cellular PM. However, it remains unclear if the ExHOS assortment is simply the result of stochasticity or if it is rooted in the observation, at distinct timepoints, of a regulated dynamical mechanism.

### A map of the exocytic machinery dynamism serves as a time ruler of exocytosis

To investigate the temporal control of ExHOS, we explored the temporal correlation of exocyst clusters with other proteins of the exocytic machinery. We first used the live-cell fluorescence intensity ratiometric assay^15^ to estimate the stoichiometry in PM-associated clusters of proteins representative of main functional steps of exocytosis: Sec2-mNeonGreen (Sec2-mNG) (vesicle transport), mNG-Sec9 (vesicle fusion) and Sec17-mNG and Sec18-mNG (protein recycling). We found that, on average, each exocytic event consists of 8 ± 1, 9 ± 1, 26 ± 4 and 37 ± 5 copies of Sec2-mNG, mNG-Sec9, Sec17-mNG and Sec18-mNG, respectively (Figures 3A and S5), which is consistent with a ∼1:1 stoichiometry between the exocyst and Sec2, Sec9, and the Sec17-Sec18 complex (which is formed by 4 copies of Sec17 and 6 copies of Sec18)^18–20^.

**Figure 3.**
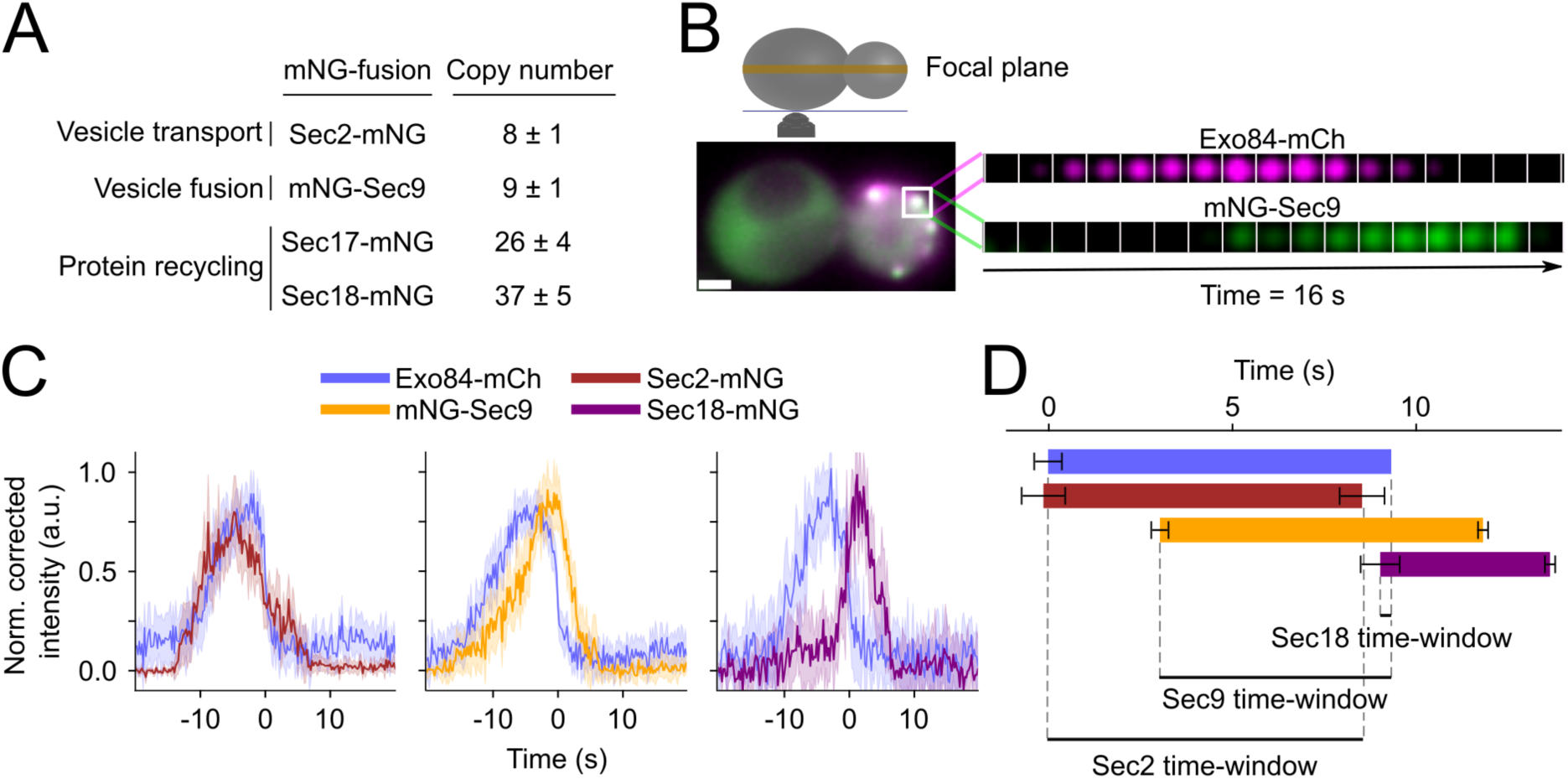
Live-cell imaging of stoichiometry and dynamics in exocytic events. (A) Average copy number of protein clusters representative of different functional steps in exocytosis (Figure S5 and Methods). Data is shown as mean ± pooled SEM. (B) Illustrative cell expressing exocyst-mCh (magenta) and mNG-Sec9 (green) simultaneously imaged at the cell equatorial plane. The insets show an illustrative track (1 s between frames for visualization). Scale bar: 1 μm. (C) Average intensity profiles of exocyst-mCh and Sec2-mNG (left, 34 tracks), mNG-Sec9 (center, 42 tracks) and Sec18-mNG (right, 14 tracks) clusters obtained by simultaneous dual-color imaging (115-120 ms/frame). The lighter areas represent 95% Confidence Intervals (95% CI). (D) Average temporal map of exocytosis derived from two-color particle tracking. Time-windows are defined as the time when Exo84-mCh colocalizes with mNG fusions. Error bars represent the pooled SEM of 3 biological replicates.

Live-cell imaging also allowed us to quantify, with 115-120-ms temporal resolution, the dynamics of these proteins in exocytic events. We performed simultaneous two-color diffraction-limited imaging at the equatorial plane of yeast cells expressing Exo84 fused to mCherry (exocyst-mCh) and a mNG-tagged marker (Sec2-mNG, mNG-Sec9 or Sec18-mNG; Figure 3B). Based on quantitative features derived from individual tracks (e.g., displacement, track quality, etc.), we set a semi-automated pipeline for the tracking of individual events of exocytosis (Methods). Particle tracking of 90 exocytic events allowed us to quantify the average dynamics of mNG-tagged markers in relation to the average intensity profile along the lifetime of exocyst-mCh clusters (Figure 3C and Methods). Under our experimental conditions, exocyst-mCh clusters present an average lifetime of 9.4 ± 0.5 s (Figure 3D). Underscoring the functional relevance of filtered tracks, more than 90% of the exocyst-mCh clusters colocalize with clusters of mNG-tagged markers (Figure S6), including the recruitment of Sec18-mNG, which we use as a proxy of the normal termination of exocytosis after vesicle fusion. In addition, 90% of the Sec2-mNG clusters immobilized to the PM coincide in time and space with the detection of immobile exocyst-mCh clusters. Overall, this demonstrates that the majority of exocyst-mCh clusters tracked by our approach correspond to functional individual events of exocytosis.

All the mNG-tagged marker tracks were aligned to the release time of the exocyst-mCh clusters, normalized by their corresponding maximum corrected intensity, and then averaged. The detection of Sec2-mNG tracks is concomitant to the clustering of exocyst-mCh, which we interpret as tethering initiation. Then, soluble SNARE mNG-Sec9 is recruited to exocytic events 3.1 ± 0.2 s after the clustering of exocyst-mCh. Sec2-mNG and mNG-Sec9 concur in the event of exocytosis for about 5.5 s. Subsequently, Sec2-mNG is released 0.8 ± 0.6 s before the release of exocyst-mCh clusters. Finally, Sec18-mNG is recruited to exocytic events 9.1 ± 0.5 s after tethering started, yielding a short, yet consistent, 0.3 s colocalization with the exocyst-mCh on average (Figure 3C and 3D). mNG-Sec9 and Sec18-mNG clusters are released from the exocytic event 2.5 ± 0.1 s and 4.3 ± 0.1 s after the release of exocyst-mCh clusters, respectively. Overall, from the biogenesis of static exocyst-mCh clusters associated with the PM to the release of Sec18-mNG, the process of exocytosis lasts 13.7 ± 0.6 s, on average. The reported temporal map indicates that the presence of different fluorescent markers defines overlapping but distinct time-windows along the exocyst-mCh cluster lifetime: Sec2 (0 s to 8.6 s), Sec9 (3.1 s to 9.4 s) and Sec18 (9.1 s to 9.4 s) (Figure 3D). These time-windows allowed us to address the temporal organization of the ExHOS.

### The ring-shaped ExHOS follows a radial expansion during exocytosis

Equipped with temporal markers, we next investigated the existence of a dynamic ring-shaped ExHOS that adjusts its radius to suit the tethering of the secretory vesicle as it approaches the PM. Unfortunately, it is not possible to continuously follow the structural dynamics of the ExHOS in the cell because of the technical limitations of SMLM. Instead, we correlated SMLM imaging of exocyst-mM with DL imaging of the temporal markers Sec2-GFP and mNG-Sec9. For each dataset, we computed the average image by aligning the clusters to their Taubin’s center (Methods). In agreement with a structurally dynamic ExHOS, exocyst-mM clusters that colocalize with Sec2-GFP present a patch-shaped average image, with a radius of 32 ± 2 nm, while those that colocalize with mNG-Sec9 present a ring-shaped average image of 37 ± 1 nm of radius (Figure 4A and Methods).

**Figure 4.**
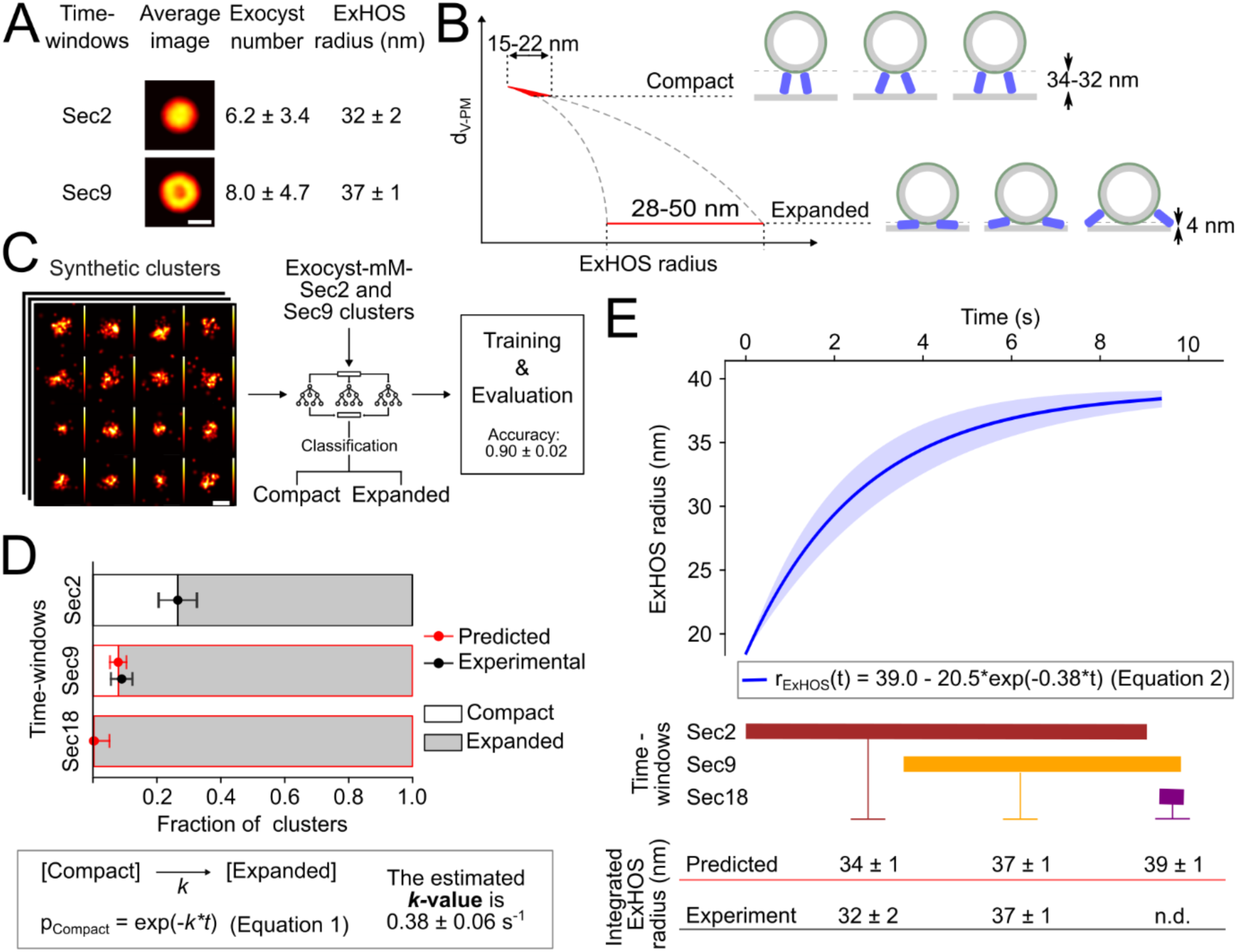
The conformational dynamics of the ExHOS time-resolved by SMLM. (A) Correlative SMLM-DL imaging of yeast cells expressing exocyst-mM and Sec2-GFP (173 clusters) or mNG-Sec9 (202 clusters). The average image, exocyst number (mean ± pooled SD, 3 biological replicates) and underlying ExHOS radius (mean ± pooled SEM, 3 biological replicates) were estimated for each time-window. Scale bar: 100 nm. (B) Plot illustrating the conformational space of modeled ExHOS for the Compact (radii from 15 to 22 nm; top) and Expanded (radii from 28 to 50 nm; bottom) classes. (C) Illustrative SMLM images of the synthetic dataset used to train a random forest classifier (Figure S7) to classify imaged exocyst-mM clusters into Compact or Expanded classes. (D) The experimental relative populations of Compact and Expanded ExHOS are plotted in black (top). The rate constant *k* for a first-order reaction was estimated for the transition from Compact to Expanded ExHOS (equation (1) and Methods) by fitting the model to the relative population of classes measured for the Sec2 time-window (bottom). The predicted relative population of classes in the Sec9 and Sec18 time-windows are shown in red (top). (E) Temporal evolution of the average ExHOS radius, *r_ExHOS_*, (equation (2) and Methods). The lighter areas represent the 95% CI of *r_ExHOS_* (top). Experimental and predicted integrated *r_ExHOS_* values for Sec2, Sec9 and Sec18 time-windows (bottom).

To estimate the underlying continuum dynamics of the ExHOS, we first discriminated between the two conformational extremes predicted by computational models. According to the modeled possible ExHOS that contain 6 to 8 exocysts (Figure 1C), ExHOS radius varies between 15 and 22 nm (defined here as the Compact ExHOS class) for the tethering at 32-34 from the PM. Instead, ExHOS radius can fluctuate between 28 nm and 50 nm (defined here as the Expanded ExHOS class) for the tethering at 4 nm from the PM (Figure 4B). We used the synthetic dataset of ring-shaped ExHOS to train a random forest classifier to systematically classify the imaged exocyst-mM-GFP clusters according to their likelihood of representing a Compact or an Expanded ExHOS. The classifier achieved 90% accuracy on the classification of synthetic SMLM images (Figure 4C, S7 and Methods).

Overall, 27 ± 3% and 73 ± 6% of the exocyst-mM clusters correlating with Sec2-GFP were classified as Compact and Expanded ExHOS, respectively (Figure 4D). In line with our models, and taking into account the ∼20 nm resolution of SMLM, the average Compact ExHOS appears as a patch-shaped cluster, with an estimated radius of 19 ± 1 nm. The average Expanded ExHOS presents a wider ring-like shape, with an estimated radius of 36 ± 1 nm (Figure S8 and Methods). However, only 9 ± 4% of the exocyst-mM clusters correlating with mNG-Sec9 were predicted to follow a Compact ExHOS, whereas 91 ± 4% of them were classified as Expanded (Figure 4D). The average image for Expanded clusters colocalizing with mNG-Sec9 presents a ring with fewer localizations in the center, which results in a significantly larger radius (38 ± 1 nm; Figure S8). We could not detect any significant difference in the number of exocysts populating each class of clusters or time-window (Figure 4A and S4). In conclusion, the combination of SMLM and fluorescent temporal markers shows that clusters of 7 exocysts on average transition from Compact to Expanded ExHOS along the process of exocytosis.

The description of the ExHOS dynamics presented until this point is based on the idea of two well-defined conformations, Compact and Expanded. This approximation of the realistic continuous tethering mechanism to two discrete conformations allowed us to question if a switch from a Compact to an Expanded ExHOS could be explained by a structural transition following first-order reaction-like kinetics:

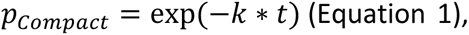

where *p_Compact_* is the relative population of the Compact class. The average rate constant *k* was estimated based on the fraction of Compact and Expanded ExHOS present in the Sec2 time-window: *k* = 0.38 ± 0.06 s^-1^ (Figure 4D and Methods). Underscoring the relevance of the model, the estimated rate constant could predict the fraction of Compact and Expanded ExHOS measured in the Sec9 time-window (Figure 4D). Consequently, this model allowed us to predict the continuous ExHOS conformational dynamics (i.e., the average ExHOS radius, *r_ExHOS_*, along the lifetime of exocyst clusters; Figure 4E):

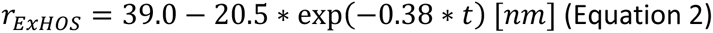

According to this model, the ExHOS initiates, on average, as a ∼19 nm radius ring and it undergoes an exponential radial expansion with a mean half-lifetime of ∼3 s. To challenge this model derived from Sec2-GFP dataset, we computed the integrated average radius for the ExHOS corresponding to Sec9 time-window. In agreement with the Equation 2, the average radius measured for the exocyst-mM clusters that correlate mNG-Sec9 (37 ± 1 nm) matches the predicted average radius (37 ± 1 nm) (Figure 4E). Remarkably, the model predicts a metastable conformation of the ExHOS with an average radius of 39 ± 1 nm 9.4 s after tethering initiation (Figure 4E). Unfortunately, the inherent short-lived colocalization between the exocyst and Sec18 precluded us from obtaining sufficient data to resolve the ExHOS during the last 300 ms of the exocyst cluster lifetime.

### Sec18 is required for exocyst recycling

To challenge the theoretical stability of the ExHOS at the final stages of exocytosis, we investigated the molecular bases that regulate the release of exocyst clusters. Interestingly, the average intensity profile shows that the release of exocyst-mCh clusters progresses in parallel to the clustering of Sec18-mNG within events of exocytosis (Figure 3C). Moreover, the correlation (r-value = 0.8) between the release of exocyst-mCh clusters and the moment when Sec18-mNG starts clustering (Figure 5A) underscores the relevance that Sec18 might have, directly or indirectly, in controlling the dynamics of the ExHOS. In contrast, neither the clustering nor the release of Sec2-mNG or mNG-Sec9 show a significant correlation with exocyst-mCh release (Figures 5A and S9). Therefore, Sec18 arises as the only tested protein that might be involved in the release and recycling of the exocyst clusters.

**Figure 5.**
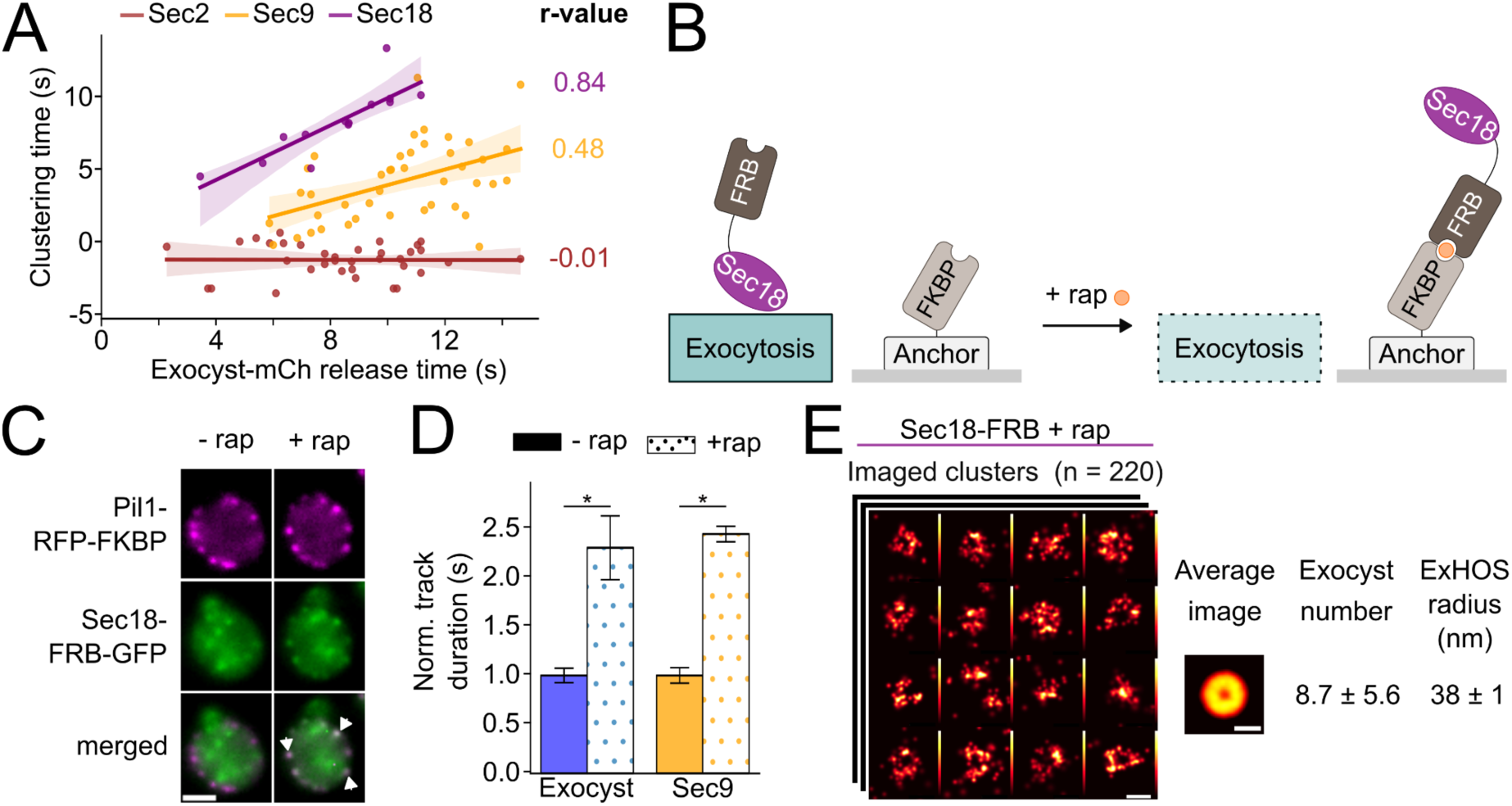
Sec18 is necessary to disassemble the ExHOS. (A) Linear models were fitted between the times for exocyst release and the clustering of Sec2 (n=34), Sec9 (n=42) and Sec18 (n=14) mNG-fusions, respectively. Lighter areas indicate 95% CI. (B) Schematic representation of the inducible Sec18 depletion. Cells were engineered to express the eisosome protein Pil1 tagged to FKBP (anchor-FKBP) and Sec18 tagged to FRB (Sec18-FRB). Upon rapamycin incubation the FKBP and FRB domains dimerize, sequestering Sec18-FRB at the anchor-FKBP. (C) Representative images of Sec18-FRB-GFP (green) recruitment to Pil1-RFP-FKBP anchors (magenta) before and after 10 min of 10 µM rapamycin incubation. Arrowheads point to anchors where Sec18-FRB-GFP has been sequestered. Scale bar: 2 μm. (D) Average lifetime of exocyst-mCh and mNG-Sec9 clusters before (n=32) and after (n=23) Sec18-FRB depletion. *=p < 0.05. (E) Illustrative exocyst-mM-mNG clusters imaged in cells depleted of Sec18-FRB (10 µM rapamycin for 10 minutes; n = 220 clusters). The average image, exocyst copy number (mean ± pooled SD, 3 biological replicates) and underlying ExHOS radius (mean ± pooled SEM, 3 biological replicates) were estimated. Scale bars: 100 nm.

To study the functional relevance of Sec18, we engineered cells for the inducible depletion of the AAA ATPase using the anchor-away approach (Figures 5B and 5C and Methods). Importantly, as the different functional steps of exocytosis are interlinked in uninterrupted cycles of tethering-fusion-recycling, impairing Sec18 function is likely to eventually impact all other stages of the process. This is especially relevant for Sec9 and the other SNAREs, as the lack of the AAA ATPase causes their sequestration on the PM^7^. To minimize the likelihood of undesired secondary functional depletions, we quantified the impact that Sec18-FRB depletion has on the recruitment of mNG-Sec9 to exocyst-mCh clusters. Upon 10 minutes of Sec18-FRB depletion, we could not detect any significant impact on the engagement of mNG-Sec9 to new cycles of exocytosis (Figure S10). Thereafter, we treated cells with rapamycin for a maximum of 10 minutes to specifically induce the depletion of Sec18-FRB.

Firstly, we measured the dynamics of exocyst-mCh and mNG-Sec9 clusters upon the depletion of Sec18-FRB. As anticipated, cells depleted for Sec18-FRB accumulate mNG-Sec9 at the PM in clusters that increase their lifetime 2.5 times (Figure 5D). In agreement with a role of the AAA ATPase in the recycling of the ExHOS, cells depleted for Sec18-FRB accumulate exocyst-mCh clusters, whose lifetime increases 2.3-fold (Figure 5D). We then explored the role of Sec18 in controlling ExHOS conformational dynamics by imaging exocyst-mM-mNG clusters by SMLM-DL upon Sec18-FRB depletion. In agreement with the continuous ExHOS conformational dynamics model (Figures 4D and 4E), nearly all the ExHOS in cells depleted for Sec18-FRB were classified as Expanded ExHOS (94 ± 4 %; Figure S8C) and the average image presents a ring-like shape with an average radius of 38 ± 1 nm (Figure 5E, S8A and S8B). Overall, our results indicate that Sec18 is required for the native disassembly of ring-shaped ExHOS during the last 300 ms of exocyst cluster lifetime, when they have a stable radius at around 38 nm, on average.

### Time-resolved ultrastructure associated with exocyst clusters: from tethering to fusion

To complement the continuous ExHOS conformational dynamics with defined functional annotations, we used cryo-CLEM. We took advantage of the fluorescent temporal markers to experimentally time-resolve the vesicle distance to the PM. We first imaged cells expressing the exocyst fused to mNG (exocyst-mNG). We targeted the acquisition of tomograms to eighteen diffraction-limited exocyst-mNG clusters. In fifteen cases, we could unambiguously correlate the presence of an isolated vesicle with the exocyst-mNG cluster (i.e., no other vesicle could be observed at a distance <250 nm). In the other three tomograms, no vesicle or PM deformations were detected (Figure 6A). We also performed cryo-CLEM on cells expressing Sec9 fused to mScarlet3 (mSc3-Sec9). We targeted the acquisition of fourteen tomograms to mSc3-Sec9 diffraction-limited clusters and in five of those tomograms we could unambiguously correlate an isolated vesicle. In the remaining nine tomograms no vesicle or PM deformations could be correlated (Figures 6B). We did not image cells expressing Sec2-mNG because this marker alone cannot distinguish tethered from non-tethered vesicles in vitrified samples. Interestingly, while exocyst-mNG-correlated vesicles display a wide spectrum of distances to the PM (i.e., from 2 to ∼37 nm), all vesicles found within the Sec9 time-window lay closer than 7 nm from the PM (Table S1).

**Figure 6.**
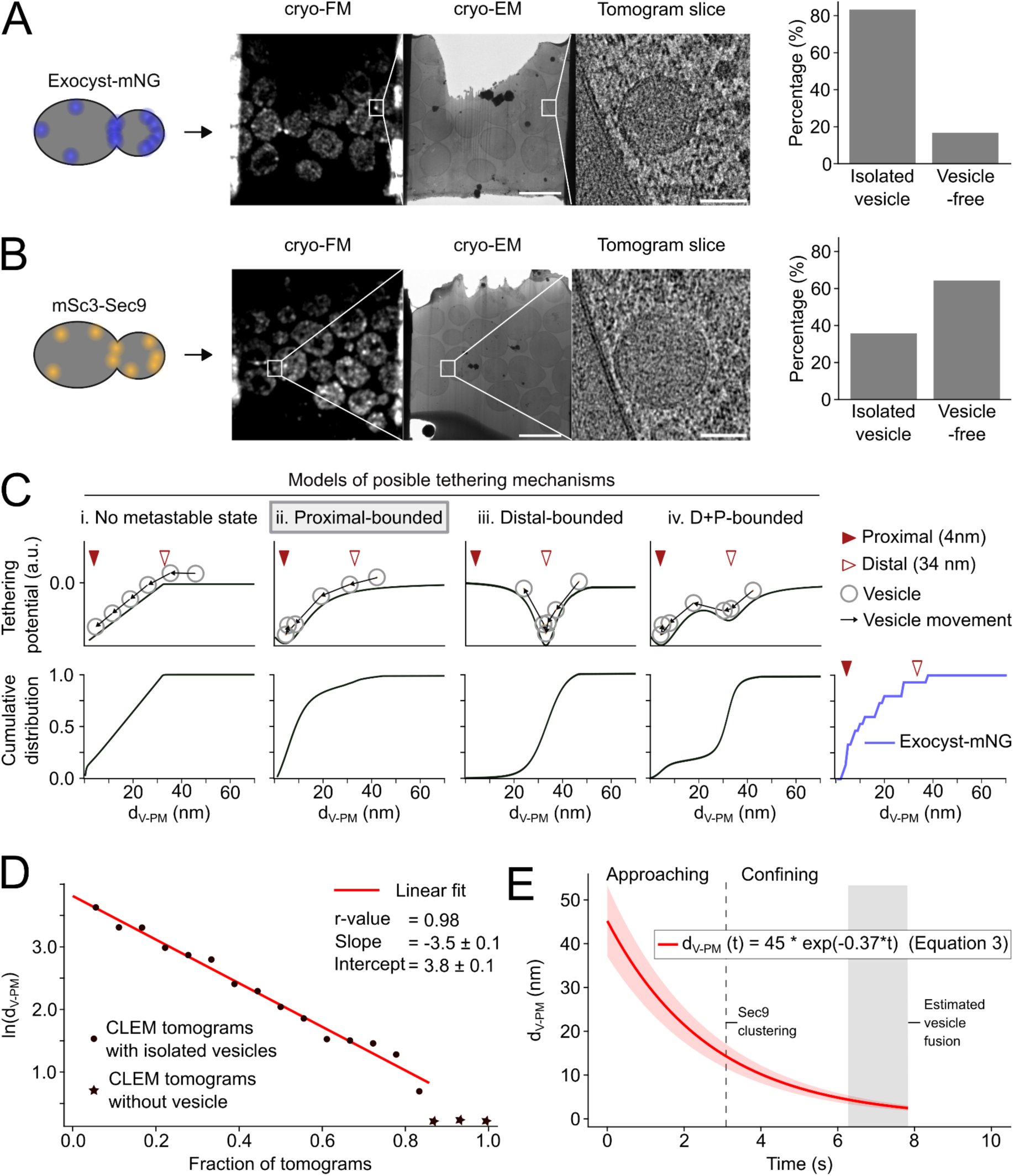
Time-resolved ultrastructure of tethering. (A-B) FIB-milled lamellae of cells expressing exocyst-mNG (A) and mSc3-Sec9 (B) were sequentially imaged by cryo-FM and cryo-TEM. Tomogram acquisition was targeted to exocyst-mNG (n=18) and mSc3-Sec9 (n=15) diffraction-limited fluorescent clusters. The fraction of diffraction-limited fluorescent spots that could, or could not, be correlated with secretory vesicles are shown (right). Scale bars: 5 μm (lamella), 50 nm (zoom-in tomogram). (C) Illustration of representative effective tethering potentials (top) and the corresponding d_V-PM_ cumulative distribution (bottom). The latter are compared with the d_V-PM_ cumulative distribution measured in tomograms targeted to exocyst-mNG in (A) (right). (D) Linear fit of the logarithm of the ordered d_V-PM_ measured in tomograms targeted to exocyst-mNG as a function of the fraction of tomograms. (E) Plotting of the continuous model of the vesicle kinetics described by Equation 3 (Methods). Lighter area indicates 95% CI by randomizing the involved parameters considering their respective errors. The Sec9 clustering time (dashed line) correlates with the transition from the Approaching to the Confining phase. The estimated time interval for vesicle fusion is indicated (gray).

Importantly, as Sec18 is known to be recruited on the exocytic *cis*-SNARE complex after vesicle fusion^8,21^, the coincidence of Sec18-mNG and exocyst-mCh suggests that the ExHOS might have a role related to vesicle fusion, in line of recently published *in vitro* reconstitutions^22^. Remarkably, in 3 out of 18 tomograms targeted to exocyst-mNG clusters (17%) and in 9 of the 14 tomograms targeted to mSc3-Sec9 clusters (64%) we could not find any nearby vesicle or PM deformation (Figures 6A and 6B). Although we could not capture fusion intermediates, our tomograms suggest that the vesicle and the PM fuse in a rapid and transient event, in agreement with experiments showing that multiple exocysts promote vesicle full fusion with the PM^23^. Hence, combining cryo-CLEM with the relative lifetimes of the exocyst-mCh and mNG-Sec9 clusters (Figures 3C and 3D), we estimated that between 6.2 s and 7.8 s after tethering initiation the vesicle fuses with the PM in the presence of the ExHOS (Methods).

Overall, the integration of SMLM-DL, genetics and cryo-CLEM implies that, as tethering progresses, the vesicle is unidirectionally dragged towards the PM. Upon fusion, the ExHOS preserves its conformation on the PM despite the lack of a vesicle that scaffolds the ring-shaped conformation.

### Modeling the kinetics of secretory vesicles reveals two functional phases

Beyond time-resolving the ultrastructure of exocytosis, cryo-CLEM can also be used to estimate the kinetics that best describes the vesicle movement along the process of tethering. Importantly, this estimation is independent of the continuous ExHOS conformational dynamics. We first computed the theoretical distribution of distances between the vesicle and the PM following tethering mechanisms with different kinetics (Figure 6C). We considered four different kinetic modalities of the vesicle approaching to the PM: i) no metastable state (i.e., diffusion-limited tethering), ii) a potential minimum that confines vesicles in a metastable Proximal tethering (in the example of the figure the confinement happens at ∼4 nm), iii) a potential minimum that confines vesicles in a metastable Distal tethering (in the example of the figure the confinement happens at ∼34 nm), and iv) a combination of both potential minima.

To experimentally determine the type of kinetics involved in secretory vesicle tethering, we employed the distribution of the closest distance to the PM for each vesicle correlated with exocyst-mNG (Figure 6C). The cumulative distribution of tethered vesicles shows that exocyst-mediated tethering follows a mechanism that features a Proximal metastable state, at about 4 nm from the PM, where vesicles are ultimately confined (Figure 6C). Furthermore, we found that the distribution of distances between exocyst-tethered vesicles and the PM (*d_V-PM_*) can be explained by kinetics following an exponential (Figures 6D and 6E):

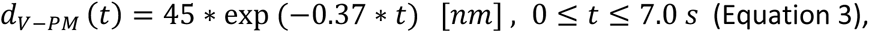

where, as derived from cryo-CLEM, fusion is estimated to take place 7s after the initiation of vesicle tethering, on average (Figure 6E and Methods). In agreement with the ultrastructural analysis done with cryo-ET (Figure 1A), this model predicts that vesicle tethering starts when the vesicle is at 45 nm from the PM (Equation 3). Tethering follows two distinguishable functional phases: the Approaching phase, where the vesicle rapidly approximates towards the PM at the beginning of tethering. And the Confining phase, where the vesicle is stabilized at about 4 nm from the PM at the culmination of tethering (Figure 6E).

### The time-resolved architecture and functional annotation of ExHOS

The quantitative characterization of fluorescent temporal markers endows the synergy between SMLM-DL and cryo-CLEM that allowed us to reconstruct the time-resolved architecture of tethering, including its continuous dynamics and functional annotation. Both the continuous ExHOS conformational dynamics (Equation 2, based on SMLM-DL) and the vesicle kinetics (Equation 3, based on cryo-CLEM) independently converge on a mechanism characterized by a fast initial dynamism that culminates in a metastable state. We combined this information with the average radius of secretory vesicles (Figure 1A), the average stoichiometry of exocyst clusters and other exocytic partners (Figures 1B and 3A), and the functional phases of tethering in an integrative model (Figure 7A and Video S1). Accordingly, tethering initiates when a cluster of 7 ± 1 exocysts arrange in a ring-shaped ExHOS with an average radius of 19 nm and an average distance of 16 nm between the centroids of adjacent exocysts. During the Approaching phase, the ExHOS radially expands while it quickly pulls the vesicle towards the PM. Concomitant to the clustering of Sec9, the energetic minimum characterizing the Confining phase implies that the ExHOS ultimately locks the vesicle at a distance of ∼4 nm from the PM. This metastable state is controlled by a stationary ExHOS with an average radius of 37 nm and an average distance of 32 nm between the centroids of adjacent exocysts. In this conformation, the exocyst is likely to coexist with the *trans*-SNARE complex (fully or partially zippered), as the distance between vesicle-associated SNAREs and SNAREs integral to the PM would allow the complex formation (Figure S11). Upon vesicle fusion with the PM, the ExHOS, with an average radius of 38 nm and 33 nm between the centroids of adjacent complexes on average, remains associated to the PM for at least 1.6 s. Eventually, the release and recycling of exocysts from the PM is completed by an unknown reaction that requires Sec18.

**Figure 7.**
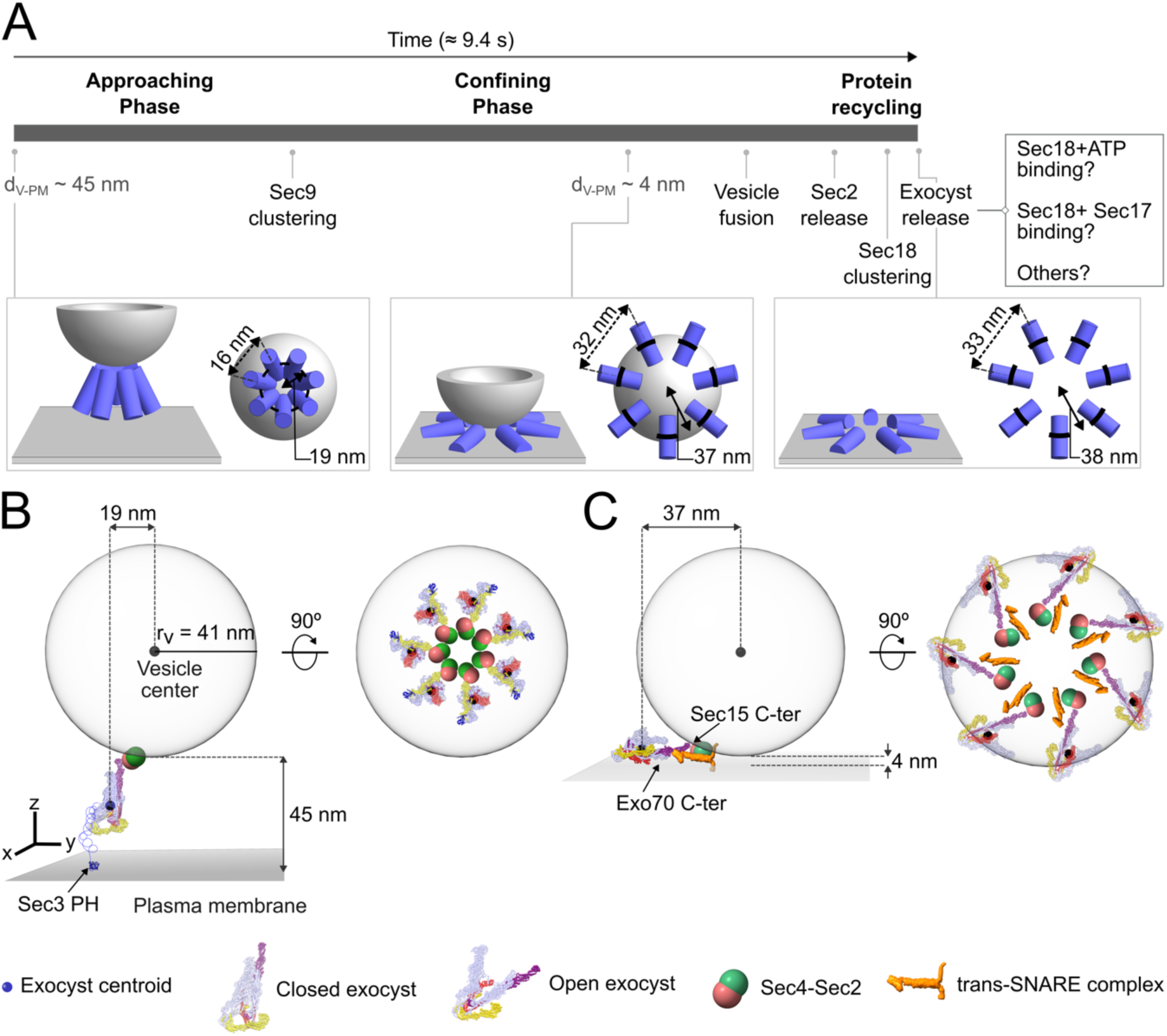
Timeline of the architecture dynamics during exocytosis. (A) Integrative model of the ExHOS and the secretory vesicle along the exocyst cluster lifetime (Video S1). The main time-points along the process are annotated in the timeline (top) with illustrations of the initial tethering (bottom-left), the time-point when the vesicle reaches 4 nm from the PM (bottom-center) and the end of the exocyst cluster lifetime (bottom-right). Possible mechanisms triggering exocyst release are outlined in the top-right box. (B-C) Speculative models illustrating the structure of the exocyst bound to a vesicle (r_v_=41 nm). The respective bottom view illustrates the arrangement of 7 exocysts. Sec3 (blue), Sec6 (yellow), Sec15 (purple) and Exo70 (red) are highlighted. (B) Beginning of tethering. The cryo-EM structure of the exocyst, between the vesicle and the PM, was positioned with its centroid at a radius of 19 nm and Sec15 C-terminus binding Sec4-Sec2^4,27^. Sec3 PH domain bound to the nearest point on the PM. (C) Confining the vesicle at 4 nm from the PM. The structure of the exocyst in an open conformation^28^ was positioned, aside to the interface between the secretory vesicle and the PM, with its center of mass at a radius of 37 nm, Sec15 C-terminus binding Sec4-Sec2 and Exo70 C-terminus binding the PM^29,30^. The *trans*-SNARE complex (orange) is located to allow insertion in the vesicle and the PM while Sec9 N-terminal is oriented to the C-terminal of Sec6^31,32^.

## DISCUSSION

Complementary imaging and modeling allowed us to resolve the native choreography of the ExHOS and cellular membranes during exocytosis.

### Regulating the stoichiometry and plasticity of the ExHOS

We showed that 7 exocysts on average organize in a ring-shaped ExHOS to control vesicle tethering. It is remarkably similar to the Munc13 hexamers that have been proposed to control mammalian synaptic exocytosis^24,25^, although the radius of synaptic vesicles is nearly half as compared to those of secretory vesicles in yeast^26^. The similarity in the stoichiometry of related –yet evolutionary distant– pathways, suggests that the number of exocysts might be critical for their function, such as the lateral immobilization of the tethered vesicle.

On the other hand, the flexible circular shape adopted by the ExHOS might have a dual role in exocytosis: i) a ring provides an ideal geometry to apply isomorphic forces that minimize the lateral movement of the vesicle while it can exert a net force orthogonal to the PM and ii) it would facilitate the interaction of the opposing membranes by allowing a higher effective contact area of the vesicle as the ExHOS expands. Simultaneously, the radial displacement of the Sec15 C-terminus during the ExHOS expansion would clear the interface between the two membranes of those proteins attached to the exocyst, such as the Sec4-Sec2 assembly (Figures 7B and 7C).

### Deriving mechanistic insight from integrative modeling

We combined the cryo-EM structure of the exocyst and the continuum architecture dynamics to reconstruct the initial moments of tethering (Figure 7B and Methods). Although tethering starts when the vesicle is at 45 nm from the PM (Figure 7B), the dimensions of the exocyst structure in isolation cannot account for such distance. Instead, our reconstruction of the initial tethering hypothesizes that the extension of the Sec3 N-terminus (residues 250 to 611 that were not resolved by cryo-EM) might be critical to determine the maximum distance at which vesicles can be tethered (Figure 7B). Indeed, this flexible stretch can link the exocyst with the Sec3 pleckstrin-homology (PH) domain that targets the complex to the PM^11^ for up to 126 nm (in an ideal full extension). Interestingly, Sec3 N-terminus nests 24 phosphorylation sites, which have been suggested to regulate Sec3 folding and binding to partners^11^. The post-translational modification of Sec3 N-terminus would provide a fast mechanism for the regulation of vesicle kinetics along the Approaching phase.

The *in vivo* architecture of the exocyst bound to a vesicle showed that Module I and Module II of the complex adopt an open conformation where they are attached by the subunit’s N-termini^13,28^. Building on this observation, we have integrated the *in vivo* architecture and the cryo-EM structure with the continuum architecture dynamics to speculate on the exocyst cluster structure when it is engaged in an ExHOS that confines secretory vesicles at 4 nm from the PM (Figure 7C). Following a 1:1:1 stoichiometry between the exocyst, the Sec4-Sec2 complex and the *trans*-SNARE complex, in this model Exo70 C-terminus could bind the PM while Sec15 C-terminus is oriented towards the Sec4-Sec2 complex and Sec6 C-terminus extends towards the *trans*-SNARE complex located at the vesicle-PM interface (Figure 7C). This conformation would allow the Sec9 N-terminal domain to bind Sec6 by mirroring the Dsl1-Sec20-Use1 binding^33^ (Figures 7C). Such interaction could support the cooperation between exocyst and SNAREs to lock the vesicle at about 4 nm from the PM, as preparation for the subsequent fusion.

### SNAREs controlling the kinetics of tethering

Indeed, the coincidence of the Confining phase with the clustering of Sec9 opens the possibility that the formation of the *trans*-SNARE complex plays a central role in tethering by locking the vesicle proximal to the PM. As Sec6 preferentially interacts with the *trans*-SNARE complex^34^, the interaction between Sec6 and Sec9 could trigger the Confining phase. Remarkably, the *trans*-SNARE complex that stands out from the membranes extends over ∼ 5 nm (Figure S11B), a length that perfectly suits the vesicle tethering for the Confining phase. Interestingly, reconstitution of vesicle tethering with the isolated human exocyst detected distal tethering (32.1 nm) but could not recapitulate proximal tethering, further supporting an essential role of the SNAREs in the confinement of secretory vesicles before fusion^35^.

Why tethering at 4 nm from the PM is sustained for about 2 s before fusion is not clear. Such a metastable tethering poses striking similarities, but also differences, with related synaptic exocytosis. Current thinking in the field is that primed synaptic vesicles are confined at <5 nm from the PM, a confinement that has been proposed to accelerate cargo secretion upon signaling^36^. However, constitutive exocytosis is intrinsically uninterrupted and such a “priming” mechanism would not take place. Our results present the possibility of cooperation between the exocyst and the *trans*-SNARE complex to provide an energetic minimum at ∼4 nm from the PM (Figures 6C and 7C). Such a metastable tethering could be necessary to support protein clearing and lipid rearrangement at the interface between the two membranes before undergoing fusion.

### A possible new role for Sec18 in exocytosis

Sec18, which unwinds the *cis*-SNARE complex, is required for the native release of exocysts (Figure 5D), coupling the recycling of both complexes. It is unlikely that exocyst release involves the hydrolysis of ATP, as the unwinding of the *cis*-SNARE complex known to involve this activity^8^ occurs 2.5 s later (Figure 3C and 3D). Instead, we hypothesize that ATP binding by the D2 domain of Sec18 or the action of Sec17 (or other unknown proteins) would provide the energy required to disassemble the ExHOS (Figure 7A). Unfortunately, our attempts to resolve this mechanism by Sec18 point mutations resulted in major defects of the exocyst subcellular distribution that prevented us from deriving meaningful data.

Overall, this study provides quantitative insights on the biophysical principles that drive tethering of secretory vesicles. Mechanistic nuances such as reversibility in the dynamics or biological heterogeneity remain unattainable because of the need of averaging. For example, while the average copy of exocyst subunits is highly homotypic, the exocyst clusters show a broad distribution of brightness and lifetime that suggest larger differences among individual events (Figure 3D and S2). The relevance of such heterogeneity remains unknown. Nonetheless, we believe that resolving how the continuum landscape of ExHOS conformations is linked to the kinetics of the tethered vesicle opens up an avenue for understanding the mechanism of the exocytic machinery as an ensemble. We forecast that the continuum dynamics of proteinaceous higher-order structures and membrane deformation kinetics will be central in the study of other pathways in membrane biology.

## METHODS

### Yeast strains

*Saccharomyces cerevisiae* strains were derived from BY4741 and BY4742 backgrounds (Invitrogen). Standard PCR-based genetic manipulation methods were used to generate strains with C-terminally tagged proteins^37^. Seamless N-terminal tagging was done as described by Khmelinskii *et al*., 2011^38^. All inserts were confirmed by PCR. Fluorescent tags were also confirmed by live-cell imaging. All strains used in this study are listed in Table S2.

### Cryo-electron tomography

#### Sample preparation and tomogram acquisition

Cells expressing exocyst-mNG or mSc3-Sec9 (Table S1) were grown in YPD at 30 °C and 200 rpm overnight. The next morning, they were diluted and grown until they reached OD_600_ = 0.6, thereafter glycerol was added to a final concentration of 5%. 4 µl of cells were deposited on glow-discharged Cu 200 mesh R1.2/1.3 grids (Quantifoil), 3.5 uL on the front and 0.5 uL at the back side. The grids were blotted and then plunged frozen into liquid ethane by back-side blotting for 4 s using an automated Plunge Freezer EM GP2 (Leica).

Grids were clipped in FIB-compatible autogrid cartridges, loaded into an Aquilos 2 Cryo-FIB (ThermoFisher Scientific) and coated with platinum. Lamellae were prepared with AutoTEM software to a target thickness of 160-180 nm. Automatic rough milling was performed with 3 steps, with 1 nA, 0.5 nA and 0.3 nA respectively. After rough milling on all sites, automatic polishing was performed with 50 pA and 30 pA. Lastly, manual polishing with 30 pA at 0.5⁰ overtilt was performed to further improve the quality of the lamellae.

Cryo-fluorescence Z-stack images of lamellae were acquired with 100X objective (0.75 NA) in AiryScan mode on LSM900 (Zeiss) equipped with a cryo-stage (Linkam). 488 nm and 561 nm lasers were used to acquire exocyst-mNG and mSc3-Sec9 images, respectively. The Z-stacks were aligned with StackReg^39^, then maximum intensity projection was done in Fiji for later correlation.

At EMBL Imaging Centre, grids with lamellae were loaded into a Titan Krios G4 (ThermoFisher Scientific) equipped with SelectrisX energy filter and Falcon4i camera, operated with SerialEM software^40^. Medium magnification montages of the lamellae were acquired and blended with Etomo^41^. Then medium-mag-montages and their corresponding fluorescence map were registered in ICY with ec-CLEMv2 plugin^42^, so that the fluorescence signal could be used to guide the high-magnification tilt series collection. Dose-symmetric tilt-series were acquired in SerialEM with PACE-tomo^43^ during three sessions with different acquisition parameters. The tilts series were recorded over a tilt range of 108° where possible, starting from the milling angle, with an angular increment of 2° or 3°. The total dose of the tilt series was 140e/A^2^. The raw frames were acquired and stored in EER format, then later processed to reduce beam-induced motion. Tilt-series were acquired with a pixel size between 1.96 and 3.04 Å, with defocus.

#### Tomographic processing

A total of 406 tilt series were acquired. After visual inspection, a total of 134 were deemed viable. On those sets, frame alignment for motion correction was performed with the implementation of the UCSF motioncor2 program in Relion 4^44^ using whole frame correction or 25 patches depending on the signal of the movies, and tomographic alignment of each tilt series was performed with IMOD 4.11.12 using patch-tracking^45^ and AreTomo 1.2.0^46^. 46 tilt series were aligned with AreTomo and reconstructed with the SART algorithm provided by the package, and 88 tilt series were aligned by IMOD and reconstructed through its SIRT-like reconstruction protocol. The 134 resulting tomograms were inspected to identify obvious misalignment problems or lack of signal, leading to the further discard of 61 tomograms.

#### Selection of isolated vesicles

To select isolated vesicles that correlated with exocyst-mNG or mSc3-Sec9 clusters, the following criteria were used: i) a diffraction-limited fluorescence spot had to be observed close to the PM, ii) the vesicle had to be closer than 70 nm from the PM, iii) the vesicle had to be at the center of the tomogram (i.e., >250 nm from the edge), and iv) no vesicles closer than 250 nm could be observed.

### Segmentation of membranes

Membrain-Seg^47^ was used to identify pixels potentially belonging to membranes using the pre-trained MemBraine_seg_v10_alpha weights on a version of the tomograms downsampled four times. For each tomogram, the normalized score map produced by Membrain-Seg was binarized (with an intensity threshold of 0). The content of the binarized map was analyzed through Matlab’s *bwconcomp* function to reveal the connected components in the tomogram.

Connected parts were divided into coherent supervoxels with an in-house, Matlab based code, aiming at identifying coherent regions of a characteristic volume of about 5000 voxels, assumed to be fragments of membranes. Edge filtering of the corresponding regions in the tomogram allowed for assigning to each supervoxel a normal vector that captured the orientation of the membrane fragment it contained.

The centroids of those supervoxels were then clustered by DBSCAN in its Matlab implementation. We used an *ad-hoc* built distance metrics that promotes grouping supervoxels according to local norm similarity and to mutual positioning in an orthogonal arrangement to both normals. In general, this automatic procedure produces very few classification errors: assignment of supervoxels from different membranes to the same cluster is a rare event. Also rare, but more frequent, is the failure to cluster all fragments of the same membrane into the same class: in some cases, the PM of a yeast cell appears not as a single entity but distributed in a few areas.

Such cases were identified and manually curated during visual inspection of results. Finally, in order to generate a smooth representation of each membrane, the centroids in a class were used as input for the alphaShape algorithm in Matlab with the smallest alpha radius for which all centroids are connected. This results in a triangulation of arbitrary mesh parameters, allowing for a fine sampling of the membrane geometry.

### Measurement of vesicle-PM distance

The simple geometry of vesicles made it possible to identify them through cross-correlation followed by fast visual curation. The distance from each vesicle to the closest point in the PM was computed by searching to the closest barycenter in the smooth triangulation provided in the previous step.

### Live-cell imaging

#### Sample preparation

For all live-cell imaging experiments, cells were grown at 30 °C and 180 rpm overnight until saturation. The next morning, cells were diluted in low fluorescence media (Yeast Nitrogen Base Low Fluorescence without Amino acids, Folic Acid and Riboflavin, 2% Complete Supplement Mixture Drop-out without Trp, and 2% glucose), grown for 5 h until OD_600_ = 1 approximately, and attached to Concanavalin A-coated coverslips.

#### Microscope setups

Images were obtained with a Nikon Eclipse Ti2-E inverted microscope configured in 3 different setups: 1) Prime 95B camera (Photometrics), 100X/1.45 NA objective (Nikon), 488 nm laser (Coherent, OBIS) with iLas2 controlled by Modular V2.0 software (GATACA) for an homogeneous oblique illumination and a filter cube containing ZET488/561x, ZET488/561m-TRF and ZT488/561rpc as excitation, emission, and dichroic filters, respectively (all from Chroma); 2) sCMOS Zyla 4.2 camera (Andor, Oxford Instruments), SR HP Apo TIRF 100x/1.49 objective (Nikon), 488 nm and 561 nm lasers (Coherent, OBIS) with iLas2 controlled by Modular V2.0 software (GATACA) and a dual bandpass filter cube (ZET488/561x, ZT488/561rpc and ZET488/561m-TRF as excitation, dichroic and emission filters, respectively), and 3) Prime 95B camera (Photometrics), SR HP Apo TIRF 100x/1.49 objective (Nikon), a SpectraX LED system (Lumencor) and a dual BP filter cube (ZET488/561x, ZT488/561rpc and ZET488/561m-TRF as excitation, dichroic and emission filters) to minimize chromatic aberrations. In all cases, the microscope was controlled by MicroManager software^48^.

#### Live-cell fluorescence ratiometric assay

Cells expressing a target protein endogenously tagged to a fluorescent protein (i.e., Sec3-GFP, Sec5-GFP, Sec6-GFP, Sec8-GFP, Sec10-GFP, Sec15-GFP, Exo70-GFP, Exo84-GFP, Sec2-mNG, mNG-Sec9, Sec17-mNG or Sec18-mNG) were mixed at a 1:1 ratio with cells expressing Cse4-GFP or Cse4-mNG and attached to a Concanavalin A-coated coverslip. Z-stacks of 21 frames separated by 250 nm were acquired using a 100 ms exposure, utilizing microscope setup 1 (see microscope setups section). The average copy number of the reference, Cse4, has been recently quantified by Single Molecule Localization Microscopy (SMLM)^16^.

Cse4 clusters in cells undergoing anaphase were tracked over the Z-stack using TrackMate^49^ with a spot diameter of 7 pixels. The plane with the highest corrected total intensity (CTI) was determined by *CTI* = *I_spot_* – *I_BG_* ∗ (*A_spot_*/*A_BG_*), where *I*_spot_/*I_BG_* and *A_spot_*_/*BG*_represent the integrated intensity and area of the selected spot and the local background, respectively. The local background is defined as the circular region with twice the diameter of the selected spot and that surrounds this spot. If CTI maximum occurred in the first or last frame, the Cse4 cluster was discarded. Afterwards, PM-associated, isolated and static target protein clusters (fluorescent spots without apparent in-plane movement) were manually chosen. Those events were tracked with a spot-diameter of 3 pixels and the CTI was calculated as explained before.

The target protein copy number (*n_tar_*) was calculated by the stoichiometry ratio *n_tar_* = *n_ref_* ∗ (*I_tar_*/*I_ref_*), where *n_ref_* is the number of Cse4 in a cluster (4.8 ± 2.4 Cse4 per kinetochore^16^, which leads to *n_ref_* = 76.8 ± 9.8 Cse4 per cluster during telophase or anaphase), and *I_tar_* and *I_ref_* are the median of the CTI values for the corresponding protein target and Cse4, respectively. The error associated to *n_tar_* was calculated as described in Picco *et al.*, 2015^15^. The average exocyst copy number was obtained by averaging the copy number of all exocyst subunits.

#### Single particle tracking

Images were acquired using the microscope setup 2 (see microscope setups section). A cage incubator mounted on the microscope (Okolab) was used to keep a constant 25 °C environment. Simultaneous acquisition was performed with an OptoSplit II Bypass image splitter (Cairn Research). A quadrant of 512×700 pixels centered in the illumination maxima of the camera sensor was used. Movies were acquired at 8.7 frames per second (fps) to a total length of 70 seconds. An exposure time of 80 ms was used, and laser power was manually optimized for each protein. Zyla camera was set to 540 mHz and binning 2. For Sec18 anchor-away (Figure 5D), movies were acquired after a 10 minute 10 µM rapamycin treatment at 2 fps during 2.5 min.

Raw movies were preprocessed to remove extracellular background using the rolling ball algorithm in Fiji^50^, with a ball radius of 60 pixels. Afterwards, the images were bleach-corrected by the *Exponential Fitting* plugin in Fiji^51^. Finally, a 3D Gaussian-filter with radius 0.5 pixels in X and Y axes and 2 pixels in time axis was applied to Exo84-mCh channel (hereafter channel 1 or C1).

C1 spots were tracked with TrackMate^49^, setting the *Difference of Gaussians* (DoG) method with a particle diameter of 3 pixels. *Simple LAP tracker* was set with a linking maximum distance of 1.5 pixels and a distance and frame gap of 2 pixels and 1 frame respectively. Tracks closer than 10 pixels from the image borders and those with a duration shorter than 10 frames were discarded. The polarized nature of exocytosis challenges the detection of isolated events. To aid the detection of isolated exocytotic events we included three new features into TrackMate: *Track_Quality_Enviroment, Track_Quality_Before* and *Track_Quality_After.* All three are computed over the 32-bits DoG-filtered movie. *Track_Quality_Enviroment* is defined as the mean intensity of the local background along the track (i.e., pixels at 2 and 3 pixels from the mean track centroid). A value close to zero of this feature indicates a uniform background signal along the track duration (i.e., no track overlapping has occurred). *Track_Quality_Before* and *Track_Quality_After* are the mean intensity of those pixels within a distance of 1.5 pixels from the mean track centroid ∼500 ms before the track start and ∼500 ms after the track end, respectively. High values of these features might indicate overlapping with neighboring tracks or a deficient tracking, since the intensity at the exocytic site is expected to be close to background levels just before and after exocyst clustering. Integrating these three features with those computed by TrackMate resulted in a list of filtered tracks, where the clustering and release of the exocyst is clearly identified. Tracks were then manually curated.

For those manually curated C1-tracks, we inspected the second channel in the same location of the respective C1-track to look for the presence of a diffraction-limited spot of the corresponding mNG-tagged protein (hereafter channel 2 or C2). If unambiguously colocalized in space with the C1-track, they were tracked using TrackMate with *ad-hoc* adjustments to optimize the C2 spot-detection and linking. Once C2 tracking was completed, clustering and release times were computed with respect to the last detected frame in C1 (TRACK_STOP feature), i.e., C2-clustering time = (TRACK_START_C2 - TRACK_STOP_C1) and C2-release time = (TRACK_STOP_C2 - TRACK_STOP_C1). In the specific case of Sec18-mNG, the corrected intensity profiles for C1 and C2, defined as (*Mean_Spot_Intensity - Mean_local_Background_Intensity*), were extracted from the raw images; where *Mean_Spot_Intensity* is the mean of intensities of pixels within 1.5 pixels from the spot centroid, and *Mean_local_Background_Intensity* is the mean of intensities of pixels located at 2 and 3 pixels from the spot centroid. The profiles of Exo84-mCh and Sec18-mNG were aligned to the TRACK_STOP_C1 frame, normalized to their corresponding maximum value and then averaged.

#### Sec18 anchor-away recruitment score test

This approach is based on the sequestration of a protein tagged to the FKBP-rapamycin binding (FRB) domain by a rapamycin-based inducible translocation to an anchoring platform labeled with the FK506-binding protein (FKBP)^52^. To induce the depletion of Sec18 from exocytic sites, we fused the FK506-binding protein (FKBP) to the C-terminus of Pil1 (Pil1-FKBP), a BAR domain containing protein that is a core component of eisosomes^53^. To be able to visualize the anchors, we labelled them with an RFP tag (Pil1-FKBP-RFP). We fused the FKBP-rapamycin binding (FRB) domain and GFP to the C-terminus of Sec18 (Sec18-FRB-GFP). FKBP and FRB tags heterodimerize in the presence of the small molecule rapamycin^52,54^. To induce the recruitment of Sec18-FRB-GFP to the Pil1-FKBP-RFP anchor, cells were attached to Concanavalin A-coated coverslips and rapamycin was added to a final concentration of 10 µM. Wide-field fluorescent microscopy images were obtained using microscope setup 3 (see microscope setups section). Red- and green-channel images were sequentially acquired every 5 min. Exposure times were 200 ms and 1000 ms for red and green channels, respectively.

The workflow for analyzing colocalization between the anchor and the target proteins was implemented as an ImageJ macro which essentially calculated the area of overlap between spots independently segmented in images from the red (anchor) and green (target) channels. A more detailed explanation of this macro is described in Pazos *et al.*, 2023^55^. By weighting the overlapping area by the total area of the green-labeled protein we obtained a colocalization score that is indicative of the amount of Sec18-FRB-GFP translocated at the anchoring platforms. Right after 5 minutes of rapamycin treatment of the engineered cells, we could detect significant sequestration of Sec18-FRB fused to GFP (Sec18-FRB-GFP) to eisosomes decorated with Pil1-FKBP fused to RFP (Pil1-FKBP-RFP) (Figures 5C and S10A).

### Single Molecule Localization Microscopy (SMLM)

#### Sample preparation

Sample preparation was performed as previously described^56^. Briefly, yeast cells were grown at 30 °C and 180 rpm overnight until saturation. The next morning, the cultures were diluted and grown at 30°C to an OD_600_ = 0.8. 2 ml of culture was pelleted (2500 rpm), resuspended in 100 µl of YPD and pipetted onto a Concanavalin A-coated coverslip. After 15 minutes in a humidified atmosphere, the non-attached cells were removed by decantation. For the Sec18 anchor-away, after 5 minutes of attachment, rapamycin was added to a final concentration of 10 µM and incubated for 10 more minutes. Coverslips were then submerged into a freshly prepared fixation solution (4% paraformaldehyde and 2% sucrose in PBS). After 15 minutes of gentle shaking, they were removed from the fixation solution and submerged into a quenching solution (100 mM NH_4_Cl in PBS) for 15 minutes with gentle shaking. Quenching was repeated twice. Finally, coverslips were washed three times in PBS for 5 min.

#### Imaging

All single-color superresolution images were acquired on a custom-built, fully automated microscope, which was built for stable long-term automated image acquisition, and featured homogeneous high-power illumination as described previously^57^. The free-space output of a commercial LightHUB laser box with 405 nm, 488 nm, 561 nm and 638 nm laser lines was focused on a speckle reducer and coupled into a multimode fiber. The output of the fiber was then imaged into the sample to homogenously illuminate a circular area of ∼1000 mm. Fluorescence was collected through a 160x NA 1.42 TIRF objective, filtered by a bandpass filter (for GFP: 525/50; for mMaple: 600/60), and focused onto an Evolve512D EMCCD camera. The z focus was optically stabilized by total internally reflecting an additional IR laser off the coverslip onto a quadrant photo diode, which was coupled into an electronic feedback loop with the piezo objective positioner. Z focus stability was typically better than 5 nm/h. All microscope components are controlled by a custom-written plugin for µManager^58^.

Samples were mounted in 1 mL of imaging buffer (50 mM Tris-HCl pH 8 in 95% D_2_O) and sealed with parafilm to prevent evaporation. The back focal plane was imaged and visually inspected to make sure no air bubbles were present in the immersion oil. Then, videos of 20000-50000 frames were acquired using the 561 nm laser at 7-9 kW/cm^2^ with an exposure time of 30 ms, and a camera EM gain and readout rate of 200 and 10 MHz, respectively. During acquisition, mMaple was slowly photoconverted to its red state by 405 nm illumination. 405 nm pulse length was controlled through an automatic feedback loop to maintain the number of blinks constant. When all mMaple was photoconverted and no more blinking was observed, acquisition was stopped. Photoconversion parameters were optimized so that no overlapping point spread functions (PSFs) were observed. After acquiring SMLM data, a diffraction-limited snapshot of GFP or mNG signal was collected using a 488 nm laser and the same camera parameters as previously described. To automatically image multiple regions of one sample, the stage position list of µManager was used to define a grid of 80-100 positions. There was a separation of 200 µm in each X and Y direction between positions to avoid undesired photoactivation and photobleaching due to scattered light. A waiting time of 3 s was established between positions to allow for mechanical equilibration of the stage after its movement.

#### Data processing and analysis

Super-resolution microscopy analysis platform (SMAP) was used to reconstruct SMLM data^59^. All images were automatically fitted during acquisition. For the reconstruction, all localizations found in consecutive frames within a range of 75 nm were grouped into one localization. Localizations with a PSF standard deviation greater than 160 nm and a localization precision that exceeded 25 nm were discarded. To visualize the images, localizations were rendered using Gaussian kernels with a standard deviation proportional to their localization precision. A minimum Gaussian standard deviation of 6 nm was used.

To correct for chromatic aberrations between SMLM and diffraction-limited (DL) images, a predetermined transformation was used to align red (photoconverted mMaple) and green (GFP or mNG) images. The transformation was created by imaging and localizing more than 1000 TetraSpeck beads in both channels. Then, a projective transformation was created from their positions. This correction was also done using SMAP.

To select isolated exocytic sites, cells were first automatically segmented by the “SegmentCellsCME” plugin from SMAP. Each cell was visualized using SMAP ROI manager, from where individual clusters (correlating to isolated DL GFP or mNG spots) were identified. Exocyst clusters were only segmented at the center of the cell surface to avoid tilted structures. Those DL spots that were not in the central region of the cell surface or that did not colocalize with mMaple signal were discarded. Finally, just those selected sites containing more than 30 localizations –which correspond to 3 exocysts on average– were considered for further analysis.

### Number of molecules by SMLM

Nuclear pore complex (NPC) proteins can be used as reference standards for assessing molecular counting in SMLM experiments^17^. Following this idea, yeast cells expressing Nup188, an NPC subunit, labelled with mMaple (Nup188-mM), were imaged using the optical settings previously described. The segmentation of Nup188-mM clusters was done manually using the *ROI Manager* interface of SMAP. The mean (*L_R_*) and standard deviation (*σ_R_*) of the distribution of localizations per Nup188-mM cluster was used to estimate the number of exocysts (*N_T_*), by computing the ratio *N_T_* = (*L_T_*/*L_R_*) ∗ *N_R_*/3, where *L_T_* is the mean number localizations per exocyst-mM or exocyst-mM-GFP cluster, *N_R_* = 16 is the average number of Nup188 per NPC^17^ and the factor 3 accounts for each exocyst being labelled with 3 mMaples. The error associated with *N_T_* is calculated as 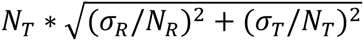.

### Modeling the possible conformational space of the ExHOS

We aimed to model the space of possible higher-order structures in which multiple exocysts can bind the vesicle and the PM simultaneously. To do this, the exocyst was represented as rigid cylinder with a height (*L_e_*) of 32 nm and a diameter (*W_e_*) of 13 nm, based on the overall exocyst dimension in the cryo-EM structure (PDB: 5YFP). Note that this is likely an underestimation of the exocyst dimensions as 18% of the exocyst amino acids, mostly Sec3 and Exo84 N-termini, could not be resolved. The vesicle was represented as a sphere of radius (*r_v_*) of 41.2 nm (this value was derived from the Cryo-ET experiments, see Figure 1). Additionally, the effective radius (*r_eff_*) of the vesicle was increased by 4 nm to account for the minimal space that the Sec4-Sec2 complex occupies on the vesicle surface (PDB: 2OCY). Assuming (i) the exocyst-vesicle and exocyst-PM binding sites are located at the extreme of the axial axis of the cylinder and (ii) the exocysts are bound radially outwards to the vesicle, the allowed configurations can be parametrized with an angle *θ* varying in the range [0, *π*/2], where *θ* = 0 when the longest axis of the exocysts are orthogonal to the PM, and *θ* = *π*/2 when the longest axis of the exocysts are parallel to the PM.

For a given exocyst, we defined the coordinates of the contact point with the PM {*x*_0_, *y*_0_, *z*_0_} and the coordinates of the contact point with Sec4-Sec2 complex at the vesicle surface {*x*_1_, *y*_1_, *z*_1_} as follows:

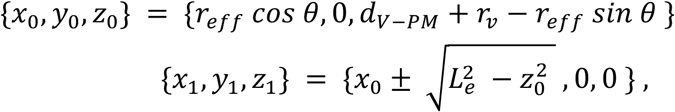

where *d_V-PM_* and 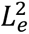 are the vesicle-PM distance and the cylinder length, respectively. We were particularly interested in the radial position (respect to the shortest axis between the vesicle and the PM) of the exocyst centroids (ρ), that can be computed as (*x*_0_+*x*_1_)/2. This parameter served us as descriptor of the ExHOS.

The underlying organization of the ExHOS highly depends on the conformational relationship among the exocysts present. For example, a heterogeneous ExHOS where the multiple exocysts are positioned in random locations, but always fulfilling the condition of binding the vesicle and the PM simultaneously. On the other hand, a homotypic ExHOS where the exocysts adopt the same conformation relative to the vesicle and the PM leads to ring-shaped ExHOSs (all ρ’s values would be equal). In this case, the maximum number of exocysts (*N_max_*_)_) can be approximated as *N_max_* = *x*_0_/*W_e_* and the coordinates for *j-*th exocyst can be obtained by applying a rotation with angle 2*π*/*N_max_* ∗ *j* around the Z-axis.

In the homotypic situation, the radius of the resulting ring-shaped ExHOS depends on the *d_V-PM_* and *N_max_*. By imposing *N_max_*≥6 (since the average number of exocysts determined by live-cell imaging was 7 ± 1 exocysts in each cluster; Figure 1B and S2), the distal tethering (*d_V-PM_*= 31 nm) leads to exocyst configurations of radius within 15-22 nm. Instead, for the proximal tethering (*d_V-PM_*= 4 nm), radii in the range 28-50 nm are possible (Figure 4B).

### Synthetic SMLM datasets

We simulated a set of SMLM images for all the possible higher-order structures modeled for distal and proximal tethering. Firstly, between 6 and 8 exocyst centroids were distributed uniformly along a ring (or randomly inside a patch) of radius *r* for a homotypic (or the heterogeneous) ExHOS. For each exocyst, three mMaple were positioned randomly, with their centroids following a two-dimensional Gaussian distribution with zero mean and a standard deviation of 10 nm along each direction. Each mMaple had a probability of emitting of 0.55, as previously determined^17^. The number of blinking events, *n*, per emitting mMaple was simulated using a log-normal distribution with parameters obtained from the fit of the number of localizations per cluster of the Nup188-mM (Figure S4). Finally, the localizations were distributed around each mMaple centroid position following a two-dimensional Gaussian distribution with zero mean and a standard deviation corresponding to an average localization precision of 20 nm along each direction. The resulting set of XY-coordinates from the generated localizations was used for image rendering and subsequent analysis. For Compact and Expanded ExHOS, synthetic images were randomly selected from *r* values in the range from 15 to 22 nm and from 28 to 50 nm, respectively.

### SMLM classification

For a given synthetic SMLM image containing the set of *N* localizations with coordinates {*x_j_*, *y_j_*}, with *j=1,···,N,* a ring center {*c_x_*, *c_y_*} and radius *R_T_* estimated using the Taubin algorithm^60^ implemented in the Python library *circle_fit* that aims to minimize the algebraic function

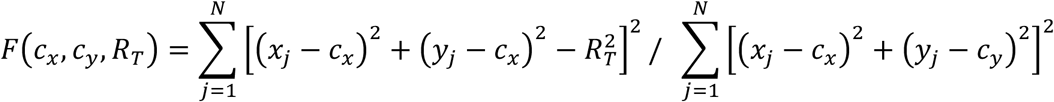

Taubin’s center estimation achieves the lower *Kanati-Cramer-Rao* bound^61^, which establishes a limit of the precision of the parameter estimation given the average SMLM resolution. In addition to *R_T_* and its associated error, a series of geometrical features were computed for all the synthetic images based on the Taubin center, including the first and second circular moment and the inner radius. The gyration radius, which does not rely on the Taubin center, was also considered. The mathematical expressions for all these geometrical features are provided in Figure S7. A matrix containing the values of the geometrical features for all the synthetic images was generated. This matrix served as input for training a random forest classifier to classify the synthetic SMLM images into the categories Compact and Expanded ExHOS. A randomized grid search was performed to fine-tune the random forest’s hyper-parameters, achieving an average accuracy of 0.90 in classifying the synthetic data. The relative importance of the geometrical features in the random forest model and the average images of the predicted classes are shown in Figure S7.

Finally, after computing the described geometrical features for the experimental set of images of isolated exocyst-mM (or exocyst-mM-GFP) clusters, the optimized random forest classifier was applied to predict the ExHOS class (Compact or Expanded). This prediction was repeated 30 times with randomizations of the training dataset, and the average prediction result was reported (Figures S8C).

### SMLM averaging and average ground-truth radius estimation

Those images belonging to a class (Compact or Expanded) were aligned to their Taubin’s center and weighted-averaged, i.e., *I_avg_*(*x*, *y*) = ∑*_i_ ω_i_ I_i_*(*x*, *y*) / ∑*_i_ ω_i_*, where *ω_i_* is the random forest-score for the *i-*th image. Then, from the average radial density *g*(*r*), the mean distance-to-center was computed: 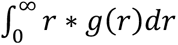. Because of the existence of estimation bias of the underlying ExHOS radius^61^, a bias-correction was implemented by creating a calibration between the underlying radii of the ExHOS and the corresponding localizations mean distance-to-center in the synthetic dataset.

### Estimation of continuous ExHOS conformational dynamics

The transition from Compact to Expanded classes of ExHOS was modeled as a first-order reaction process, according to an effective rate *k*. The relative population of the Compact ExHOS at a given time *t* after exocyst clustering, *p_Compact_* (*t*), is governed by the differential equation:

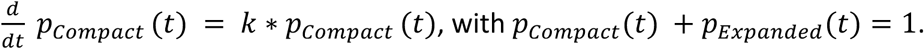

Based on the trend observed in the Sec2-GFP and mNG-Sec9 correlative SMLM experiments (Figure 4A), it is reasonable to assume that exocysts are organized exclusively in a Compact structure during the initial time-point of the tethering, leading to the condition *p_Compact_*(0) = 1. Therefore, the solution for *p_Compact_* is given by *p_Compact_* = *exp* (−*k* ∗ *t*) (Equation 1). The rate constant *k* can be estimated by solving the equation 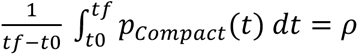, where *t*_0_and *t_f_* are the initial and final time of the given time-window, respectively, and *ρ* is the frequency of Compact ExHOS experimentally observed within this time frame. The reported errors of *k* and predicted frequency of ExHOS classes were calculated using an error-propagation scheme based on the previous equation.

Furthermore, the temporal evolution of the average ExHOS radius (*r_ExHOS_*) along the process of exocytosis was modeled by using the heuristic equation: *r_ExHOS_* (*t*) = *Mean*_*Compact*_*Radius* ∗ *p_Compact_*(*t*) + *Mean*_*Expanded*_*Radius* ∗ *p_Expanded_*(*t*), which leads to the Equation 2: *r_ExHOS_* = 39.0 − 20.5 ∗ exp(−0.38 ∗ *t*).

The estimated *r_ExHOS_* values for the Sec2 and Sec9 time-windows were computed as the mean value of *r_ExHOS_* (*t*) within the corresponding time intervals, i.e, 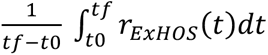.

### Estimation of the vesicle kinetics

Assuming the images obtained by cryo-CLEM provide a uniform sampling of the exocytic event lifetime and that the likelihood that multiple exocysts cluster before the arrival of the vesicle is negligible (which is supported by the enrichment of tomograms without vesicle or PM deformation in the cryo-CLEM of cells expressing mSc3-Sec9), it is reasonable to estimate the fusion time based on the frequency of exocyst-mNG correlated-tomograms with and without vesicles. Then, the 3/18 (16.7 %) tomograms that do not contain vesicles or PM deformations represent 1.6 s of the exocyst clusters lifetime (9.4 s). A similar analysis of the cryo-CLEM of cells expressing mSc3-Sec9 estimated the fusion time to be 3.2 seconds after Sec9 clustering. By combining both findings, the fusion was estimated to occur between 6.2 to 7.8 s after tethering initiation (i.e., exocyst clustering).

Additionally, we observed that the x*-*th quantile (x = 1/18,···,15/18) of the exocyst-mNG cryo-CLEM distances can be fitted with an exponential function of the form *A* ∗ exp (−*x*/*b*) (Figure 6D). Thus, we can assume that parameters *A* would be the longest distance at which a vesicle can be tethered by the exocyst to the PM and 1/*k_z_* the relative time *t*_F_/τ, with *τ* being a characteristic time of the vesicle tethering dynamics and *t*_H_the total sampling time (9.4 s). Combining these results, we determined that the motion that drags the vesicle towards the PM during tethering is dictated by the temporal evolution of the vesicle-PM distance described by Equation 3:

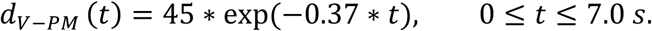

### Modeling the yeast *cis*-SNARE complex

The structure of the yeast *cis*-SNARE complex was predicted with AlphaFold3 Server^62^ using the fasta sequences of Sec9 (Uniprot ID P40357), Sso1 (Uniprot ID P32867) and Snc1 (Uniprot ID P31109) proteins. The top ranked prediction was used for analysis of the AlphaFold3 metrics^62,63^ and posterior visualization. Only the confident regions (pLDDT >70) were taken into account to represent the ribbon silhouette of the *cis*-SNARE complex (Figure S11A).

### Modeling the exocytic trans-SNARE complex

Modeling of the *trans*-SNARE complex conformation was based on the mammalian *trans*-SNARE complex simulated previously^64^. To identify the juxtamembrane linker (L) and the transmembrane region (TMR) of Sso1 and Snc1 proteins we performed a multiple sequence alignment with the mammalian homologs Syntaxin1 (Uniprot ID P32851) and Synaptobrevin2 (Uniprot ID P63045) (Figure S11B). The multiple sequence alignment was performed with Clustal Omega^65^. We used MODELLER^66^ to predict, by comparative modeling, the yeast *trans*-SNARE complex using the simulated mammalian *trans*-SNARE complex as template structure. Finally, we used ChimeraX^67^ to represent the yeast *trans*-SNARE complex with a density map of 10 Å resolution.

## ACKNOWLEDGEMENTS

We thank Francesc Posas, Manuel Mendoza, Michael Knopp and Michal Skruzny for expert help and the sharing of plasmids and reagents. We acknowledge the access and services provided by the Imaging Centre at the European Molecular Biology Laboratory (EMBL IC) and Zhengyi Yang for the technical support, generously supported by the Boehringer Ingelheim Foundation. This work was supported by iNEXT-Discovery funding (PID: 24954), the Human Frontiers Science Program (HFSP) (grant RGP0017/2020) and by Spanish funding agency [PID2021-127773NB-I00 and PID2021-127309NB-I00 funded by MICIU/AEI/10.13039/501100011033/FEDER/UE, the Unidad de Excelencia Maria de Maeztu [CEX2018-000792-M] and CNS2022-135349 funded by MICIU/AEI/10.13039/501100011033). We thank Alma Vivas Lago for inspection of raw electron microscopy data and Sebastien Tosi and Andrea Picco for helpful discussion on data analysis.

## RESOURCE AVAILABILITY

### Correspondence contact

Further information and requests for resources and reagents should be directed to and will be fulfilled by Oriol Gallego.

### Materials availability

All strains generated in this study are available upon request.

### Data and code availability

All original code and raw data will be available at https://github.com/GallegoLab and it will be deposited at online repositories before publication.

## AUTHOR CONTRIBUTIONS

Conceptualization, M.P.-T., S.O. and O.G.; methodology, M.P.-T., S.O., A.M., J.R., D.C.-D., C.M. and O.G.; software, S.O., E.K. and D.C.-D.; formal analysis, S.O., R.C. and C.M.; investigation, M.P.-T., S.M., A.C., L.B., P.H. and M.M.; resources, M.I.-S, B.O., J.R., D.C.-D., C.M. and O.G., writing – original draft, M.P.-T., S.O. and O.G.; writing – review & editing, M.P.-T, S.O., S.M., R.C., A.C.H., L.B., P.H., J.R., D.C.-D., C.M. and O.G.; visualization, M.P.-T., S.O. and O.G.; supervision, D.C.-D, C.M. and O.G.; project administration O.G.; funding acquisition, A.M., J.R., D.C.D and O.G.

## DECLARATION OF INTERESTS

The authors declare no competing interests.

## SUPPLEMENTAL INFORMATION

**Document S1. Figures S1–S11 and Tables S1-S2**

**Video S1. Integrative modeling of the architecture dynamics controlling tethering of secretory vesicles**

The temporal map and the functions describing the ExHOS radius dynamics as well as the vesicle kinetics are shown (top). The gray bar indicates the time progression during exocytosis. The timer (center) indicates the time after tethering initiation. Time-lapsed video illustrating the architecture dynamics that control the tethering of a secretory vesicle (side view, bottom-center; and bottom view, bottom-right). The average stoichiometry for the different proteins is indicated (bottom-left). The frame rate is adjusted to optimize the visualization of the key events.

## Notes

### Competing Interest Statement

The authors have declared no competing interest.

## REFERENCES

1. He, B., and Guo, W. (2009). The exocyst complex in polarized exocytosis. Curr Opin Cell Biol 21, 537–542. 10.1016/j.ceb.2009.04.007.

2. Munson, M., and Novick, P. (2006). The exocyst defrocked, a framework of rods revealed. Nat Struct Mol Biol 13, 577–581. 10.1038/nsmb1097.

3. Rossi, G., Lepore, D., Kenner, L., Czuchra, A.B., Plooster, M., Frost, A., Munson, M., and Brennwald, P. (2020). Exocyst structural changes associated with activation of tethering downstream of Rho/Cdc42 GTPases. J Cell Biol 219, e201904161. 10.1083/jcb.201904161.

4. Guo, W., Roth, D., Walch-Solimena, C., and Novick, P. (1999). The exocyst is an effector for Sec4p, targeting secretory vesicles to sites of exocytosis. EMBO J 18, 1071–1080. 10.1093/EMBOJ/18.4.1071.

5. Novick, P. (2016). Regulation of membrane traffic by Rab GEF and GAP cascades. Small GTPases 7, 252–256. 10.1080/21541248.2016.1213781.

6. Sivaram, M.V.S., Saporita, J.A., Furgason, M.L.M., Boettcher, A.J., and Munson, M. (2005). Dimerization of the exocyst protein Sec6p and its interaction with the t-SNARE Sec9p. Biochemistry 44, 6302–6311. 10.1021/BI048008Z.

7. Grote, E., Carr, C.M., and Novick, P.J. (2000). Ordering the Final Events in Yeast Exocytosis. J Cell Biol 151, 439–451. 10.1083/jcb.151.2.439.

8. Söllner, T., Bennett, M.K., Whiteheart, S.W., Scheller, R.H., and Rothman, J.E. (1993). A Protein Assembly-Disassembly Pathway In Vitro That May Correspond to Sequential Steps of Synaptic Vesicle Docking, Activation, and Fusion. Cell 75, 409–418. 10.1016/0092-8674(93)90376-2.

9. Heider, M.R., Gu, M., Duffy, C.M., Mirza, A.M., Marcotte, L.L., Walls, A.C., Farrall, N., Hakhverdyan, Z., Field, M.C., Rout, M.P., et al. (2016). Subunit connectivity, assembly determinants and architecture of the yeast exocyst complex. Nat Struct Mol Biol 23, 59–66. 10.1038/nsmb.3146.

10. Mei, K., Li, Y., Wang, S., Shao, G., Wang, J., Ding, Y., Luo, G., Yue, P., Liu, J.J., Wang, X., et al. (2018). Cryo-EM structure of the exocyst complex. Nat Struct Mol Biol 25, 139–146. 10.1038/s41594-017-0016-2.

11. Ganesan, S.J., Feyder, M.J., Chemmama, I.E., Fang, F., Rout, M.P., Chait, B.T., Shi, Y., Munson, M., and Sali, A. (2020). Integrative structure and function of the yeast exocyst complex. Protein Science 29, 1486–1501. 10.1002/PRO.3863.

12. Ahmed, S.M., Nishida-Fukuda, H., Li, Y., McDonald, W.H., Gradinaru, C.C., and Macara, I.G. (2018). Exocyst dynamics during vesicle tethering and fusion. Nat Commun 9, 5140. 10.1038/s41467-018-07467-5.

13. Picco, A., Irastorza-Azcarate, I., Specht, T., Böke, D., Pazos, I., Rivier-Cordey, A.-S., Devos, D.P., Kaksonen, M., and Gallego Correspondence, O. (2017). The In Vivo Architecture of the Exocyst Provides Structural Basis for Exocytosis. Cell 168, 400–412.e18. 10.1016/j.cell.2017.01.004.

14. Gingras, R.M., Sulpizio, A.M., Park, J., and Bretscher, A. (2022). High-resolution secretory timeline from vesicle formation at the Golgi to fusion at the plasma membrane in S. cerevisiae. Elife 11, e78750. 10.7554/ELIFE.78750.

15. Picco, A., Mund, M., Ries, J., Nédélec, F., and Kaksonen, M. (2015). Visualizing the functional architecture of the endocytic machinery. Elife 4, e04535. 10.7554/eLife.04535.001.

16. Cieslinski, K., Wu, Y.-L., Nechyporenko, L., Janice Hörner, S., Conti, D., Skruzny, M., and Ries, J. (2023). Nanoscale structural organization and stoichiometry of the budding yeast kinetochore. J Cell Biol 222, e202209094. 10.1083/jcb.202209094.

17. Thevathasan, J.V., Kahnwald, M., Cieśliński, K., Hoess, P., Peneti, S.K., Reitberger, M., Heid, D., Kasuba, K.C., Hoerner, S.J., Li, Y., et al. (2019). Nuclear pores as versatile reference standards for quantitative superresolution microscopy. Nat Methods 16, 1045–1053. 10.1038/s41592-019-0574-9.

18. Orr, A., and Wickner, W. (2022). Sec18 supports membrane fusion by promoting Sec17 membrane association. Mol Biol Cell 33, ar127. 10.1091/mbc.E22-07-0274.

19. White, I., Zhao, M., Choi, U.B., Pfuetzner, R.A., and Brunger, A.T. (2018). Structural principles of SNARE complex recognition by the AAA+ protein NSF. Elife 7, e38888. 10.7554/eLife.38888.001.

20. Zhao, M., Wu, S., Zhou, Q., Vivona, S., Cipriano, D.J., Cheng, Y., and Brunger, A.T. (2015). Mechanistic insights into the recycling machine of the SNARE complex. Nature 518, 61–67. 10.1038/nature14148.

21. Mayer, A., Wickner, W., and Haas, A. (1996). Sec18p (NSF)-Driven Release of Sec17p (alpha-SNAP) Can Precede Docking and Fusion of Yeast Vacuoles. Cell 85, 83–94. 10.1016/s0092-8674(00)81084-3.

22. Lee, C., Lepore, D., Lee, S.H., Kim, T.G., Buwa, N., Lee, J., Munson, M., and Yoon, T.Y. (2024). Exocyst stimulates multiple steps of exocytic SNARE complex assembly and vesicle fusion. Nat Struct Mol Biol. 10.1038/s41594-024-01388-2.

23. An, S.J., Rivera-Molina, F., Anneken, A., Xi, Z., McNellis, B., Polejaev, V.I., and Toomre, D. (2021). An active tethering mechanism controls the fate of vesicles. Nat Commun 12. 10.1038/s41467-021-25465-y.

24. Bera, M., Grushin, K., Sundaram, R.V.K., Shahanoor, Z., Chatterjee, A., Radhakrishnan, A., Lee, S., Padmanarayana, M., Coleman, J., Pincet, F., et al. (2023). Two successive oligomeric Munc13 assemblies scaffold vesicle docking and SNARE assembly to support neurotransmitter release. Preprint at bioRxiv. 10.1101/2023.07.14.549017.

25. Rothman, J.E., Grushin, K., Bera, M., and Pincet, F. (2023). Turbocharging synaptic transmission. FEBS Lett 597, 2233–2249. 10.1002/1873-3468.14718.

26. Zhang, B., Koh, Y.H., Beckstead, R.B., Budnik, V., Ganetzky, B., and Bellen, H.J. (1998). Synaptic vesicle size and number are regulated by a clathrin adaptor protein required for endocytosis. Neuron 21, 1465–1475. 10.1016/s0896-6273(00)80664-9.

27. Medkova, M., France, Y.E., Coleman, J., and Novick, P. (2006). The rab Exchange Factor Sec2p Reversibly Associates with the Exocyst. Mol Biol Cell 17, 2757–2769. 10.1091/mbc.E05-10.

28. Irastorza-Azcarate, I., Castañ O-Díez, D., Devos, D.P., and Gallego, O. (2019). Live-Cell Structural Biology to Solve Biological Mechanisms: The Case of the Exocyst. Structure 27, 886–892. 10.1016/j.str.2019.04.010.

29. He, B., Xi, F., Zhang, X., Zhang, J., and Guo, W. (2007). Exo70 interacts with phospholipids and mediates the targeting of the exocyst to the plasma membrane. EMBO J 26, 4053–4065. 10.1038/sj.emboj.7601834.

30. Liu, J., Zuo, X., Yue, P., and Guo, W. (2007). Phosphatidylinositol 4,5-bisphosphate mediates the targeting of the exocyst to the plasma membrane for exocytosis in mammalian cells. Mol Biol Cell 18, 4483–4492. 10.1091/mbc.E07-05-0461.

31. Shen, D., Yuan, H., Hutagalung, A., Verma, A., Kümmel, D., Wu, X., Reinisch, K., McNew, J.A., and Novick, P. (2013). The synaptobrevin homologue Snc2p recruits the exocyst to secretory vesicles by binding to Sec6p. J Cell Biol 202, 509–526. 10.1083/JCB.201211148.

32. Sivaram, M.V.S., Furgason, M.L.M., Brewer, D.N., and Munson, M. (2006). The structure of the exocyst subunit Sec6p defines a conserved architecture with diverse roles. Nature Structural & Molecular Biology 2006 13:6 13, 555–556. 10.1038/nsmb1096.

33. DAmico, K.A., Stanton, A.E., Shirkey, J.D., Travis, S.M., Jeffrey, P.D., and Hughson, F.M. (2024). Structure of a membrane tethering complex incorporating multiple SNAREs. Nat Struct Mol Biol 31, 246–254. 10.1038/s41594-023-01164-8.

34. Dubuke, M.L., Maniatis, S., Shaffer, S.A., and Munson, M. (2015). The exocyst subunit Sec6 interacts with assembled exocytic SNARE complexes. Journal of Biological Chemistry 290, 28245–28256. 10.1074/jbc.M115.673806.

35. Maib, H., and Murray, D.H. (2022). A mechanism for exocyst-mediated tethering via Arf6 and PIP5K1C-driven phosphoinositide conversion. Current Biology 32, 2821–2833.e6. 10.1016/j.cub.2022.04.089.

36. Imig, C., Min, S.W., Krinner, S., Arancillo, M., Rosenmund, C., Südhof, T.C., Rhee, J.S., Brose, N., and Cooper, B.H. (2014). The Morphological and Molecular Nature of Synaptic Vesicle Priming at Presynaptic Active Zones. Neuron 84, 416–431. 10.1016/j.neuron.2014.10.009.

37. Janke, C., Magiera, M.M., Rathfelder, N., Taxis, C., Reber, S., Maekawa, H., Moreno-Borchart, A., Doenges, G., Schwob, E., Schiebel, E., et al. (2004). A versatile toolbox for PCR-based tagging of yeast genes: new fluorescent proteins, more markers and promoter substitution cassettes. Yeast 21, 947–962. 10.1002/YEA.1142.

38. Khmelinskii, A., Meurer, M., Duishoev, N., Delhomme, N., and Knop, M. (2011). Seamless gene tagging by endonuclease-driven homologous recombination. PLoS One 6, e23794. 10.1371/JOURNAL.PONE.0023794.

39. Thévenaz, P., Ruttimann, U.E., and Unser, M. (1998). A pyramid approach to subpixel registration based on intensity. IEEE Transactions on Image Processing 7, 27–41. 10.1109/83.650848.

40. Mastronarde, D.N. (2005). Automated electron microscope tomography using robust prediction of specimen movements. J Struct Biol 152, 36–51. 10.1016/J.JSB.2005.07.007.

41. Kremer, J.R., Mastronarde, D.N., and McIntosh, J.R. (1996). Computer Visualization of Three-Dimensional Image Data Using IMOD. J Struct Biol 116, 71–76. 10.1006/JSBI.1996.0013.

42. Paul-Gilloteaux, P., Heiligenstein, X., Belle, M., Domart, M.C., Larijani, B., Collinson, L., Raposo, G., and Salamero, J. (2017). EC-CLEM: Flexible multidimensional registration software for correlative microscopies. Nat Methods 14, 102–103. 10.1038/nmeth.4170.

43. Eisenstein, F., Yanagisawa, H., Kashihara, H., Kikkawa, M., Tsukita, S., and Danev, R. (2023). Parallel cryo electron tomography on in situ lamellae. Nat Methods 20, 131–138. 10.1038/s41592-022-01690-1.

44. Scheres, S.H.W. (2012). RELION: Implementation of a Bayesian approach to cryo-EM structure determination. J Struct Biol 180, 519–530. 10.1016/J.JSB.2012.09.006.

45. Mastronarde, D.N., and Held, S.R. (2017). Automated tilt series alignment and tomographic reconstruction in IMOD. J Struct Biol 197, 102–113. 10.1016/J.JSB.2016.07.011.

46. Zheng, S., Wolff, G., Greenan, G., Chen, Z., Faas, F.G.A., Bárcena, M., Koster, A.J., Cheng, Y., and Agard, D.A. (2022). AreTomo: An integrated software package for automated marker-free, motion-corrected cryo-electron tomographic alignment and reconstruction. J Struct Biol X 6, 100068. 10.1016/J.YJSBX.2022.100068.

47. Lamm, L., Zufferey, S., Righetto, R.D., Wietrzynski, W., Yamauchi, K.A., Burt, A., Liu, Y., Zhang, H., Martinez-Sanchez, A., Ziegler, S., et al. (2024). MemBrain v2: an end-to-end tool for the analysis of membranes in cryo-electron tomography. Preprint at bioRxiv. 10.1101/2024.01.05.574336.

48. Edelstein, A.D., Tsuchida, M.A., Amodaj, N., Pinkard, H., Vale, R.D., and Stuurman, N. (2014). Advanced methods of microscope control using μManager software. J Biol Methods 1, e10. 10.14440/jbm.2014.36.

49. Tinevez, J.Y., Perry, N., Schindelin, J., Hoopes, G.M., Reynolds, G.D., Laplantine, E., Bednarek, S.Y., Shorte, S.L., and Eliceiri, K.W. (2017). TrackMate: An open and extensible platform for single-particle tracking. Methods 115, 80–90. 10.1016/J.YMETH.2016.09.016.

50. Schindelin, J., Arganda-Carreras, I., Frise, E., Kaynig, V., Longair, M., Pietzsch, T., Preibisch, S., Rueden, C., Saalfeld, S., Schmid, B., et al. (2012). Fiji: an open-source platform for biological-image analysis. Nat Methods 9, 676–682. 10.1038/nmeth.2019.

51. Miura, K. (2020). Bleach correction ImageJ plugin for compensating the photobleaching of time-lapse sequences. F1000Res 9, 1494. 10.12688/f1000research.27171.1.

52. Haruki, H., Nishikawa, J., and Laemmli, U.K. (2008). The Anchor-Away Technique: Rapid, Conditional Establishment of Yeast Mutant Phenotypes. Mol Cell 31, 925–932. 10.1016/j.molcel.2008.07.020.

53. Moreira, K.E., Walther, T.C., Aguilar, P.S., and Walter, P. (2009). Pil1 Controls Eisosome Biogenesis. Mol Biol Cell 20, 809–818. 10.1091/mbc.E08.

54. Chen, J., Zheng, X.F., Brown, E.J., and Schreiber, S.L. (1995). Identification of an 11-kDa FKBP12-rapamycin-binding domain within the 289-kDa FKBP12-rapamycin-associated protein and characterization of a critical serine residue. Proc Natl Acad Sci U S A 92, 4947–4951. 10.1073/PNAS.92.11.4947.

55. Pazos, I., Puig-Tintó, M., Betancur, L., Cordero, J., Jiménez-Menéndez, N., Abella, M., Hernández, A.C., Duran, A.G., Adachi-Fernández, E., Belmonte-Mateos, C., et al. (2023). The P4-ATPase Drs2 interacts with and stabilizes the multisubunit tethering complex TRAPPIII in yeast. EMBO Rep 24, e56134. 10.15252/embr.202256134.

56. Mund, M., van der Beek, J.A., Deschamps, J., Dmitrieff, S., Hoess, P., Monster, J.L., Picco, A., Nédélec, F., Kaksonen, M., and Ries, J. (2018). Systematic Nanoscale Analysis of Endocytosis Links Efficient Vesicle Formation to Patterned Actin Nucleation. Cell 174, 884–896.e17. 10.1016/j.cell.2018.06.032.

57. Deschamps, J., Rowald, A., and Ries, J. (2016). Efficient homogeneous illumination and optical sectioning for quantitative single-molecule localization microscopy. Opt Express 24, 28080. 10.1364/oe.24.028080.

58. Deschamps, J., and Ries, J. (2020). EMU: reconfigurable graphical user interfaces for Micro-Manager. BMC Bioinformatics 21. 10.1186/s12859-020-03727-8.

59. Ries, J. (2020). SMAP: a modular super-resolution microscopy analysis platform for SMLM data. Nat Methods 17, 870–872. 10.1038/s41592-020-0938-1.

60. Taubin, G. (1991). Estimation of Planar Curves, Surfaces, and Nonplanar Space Curves Defined by Implicit Equations with Applications to Edge and Range Image Segmentation. IEEE Trans Pattern Anal Mach Intell 13, 1115. 10.1109/34.103273.

61. Chernov, N., and Lesort, C. (2004). Statistical efficiency of curve fitting algorithms. Comput Stat Data Anal 47, 713–728. 10.1016/J.CSDA.2003.11.008.

62. Abramson, J., Adler, J., Dunger, J., Evans, R., Green, T., Pritzel, A., Ronneberger, O., Willmore, L., Ballard, A.J., Bambrick, J., et al. (2024). Accurate structure prediction of biomolecular interactions with AlphaFold 3. Nature 630, 493–500. 10.1038/s41586-024-07487-w.

63. Jumper, J., Evans, R., Pritzel, A., Green, T., Figurnov, M., Ronneberger, O., Tunyasuvunakool, K., Bates, R., Žídek, A., Potapenko, A., et al. (2021). Highly accurate protein structure prediction with AlphaFold. Nature 596, 583–589. 10.1038/s41586-021-03819-2.

64. Rizo, J., Sari, L., Jaczynska, K., Rosenmund, C., and Lin, M.M. (2024). Molecular mechanism underlying SNARE-mediated membrane fusion enlightened by all-atom molecular dynamics simulations. Proc Natl Acad Sci U S A 121, e2321447121. 10.1073/pnas.2321447121.

65. Madeira, F., Madhusoodanan, N., Lee, J., Eusebi, A., Niewielska, A., Tiv, A.R., Lopez, R., and Butcher, S. (2024). The EMBL-EBI Job Dispatcher sequence analysis tools framework in 2024. Nucleic Acids Res 52, 521–525. 10.1093/nar/gkae241.

66. Šali, A., and Blundell, T.L. (1993). Comparative Protein Modelling by Satisfaction of Spatial Restraints. J Mol Biol 234, 779–815. 10.1006/JMBI.1993.1626.

67. Meng, E.C., Goddard, T.D., Pettersen, E.F., Couch, G.S., Pearson, Z.J., Morris, J.H., and Ferrin, T.E. (2023). UCSF ChimeraX: Tools for structure building and analysis. Protein Science 32, e4792. 10.1002/PRO.4792.

