## Supplementary material for "Continuum architecture dynamics of vesicle tethering in exocytosis": Suppl_Puig-Tinto_2025.pdf

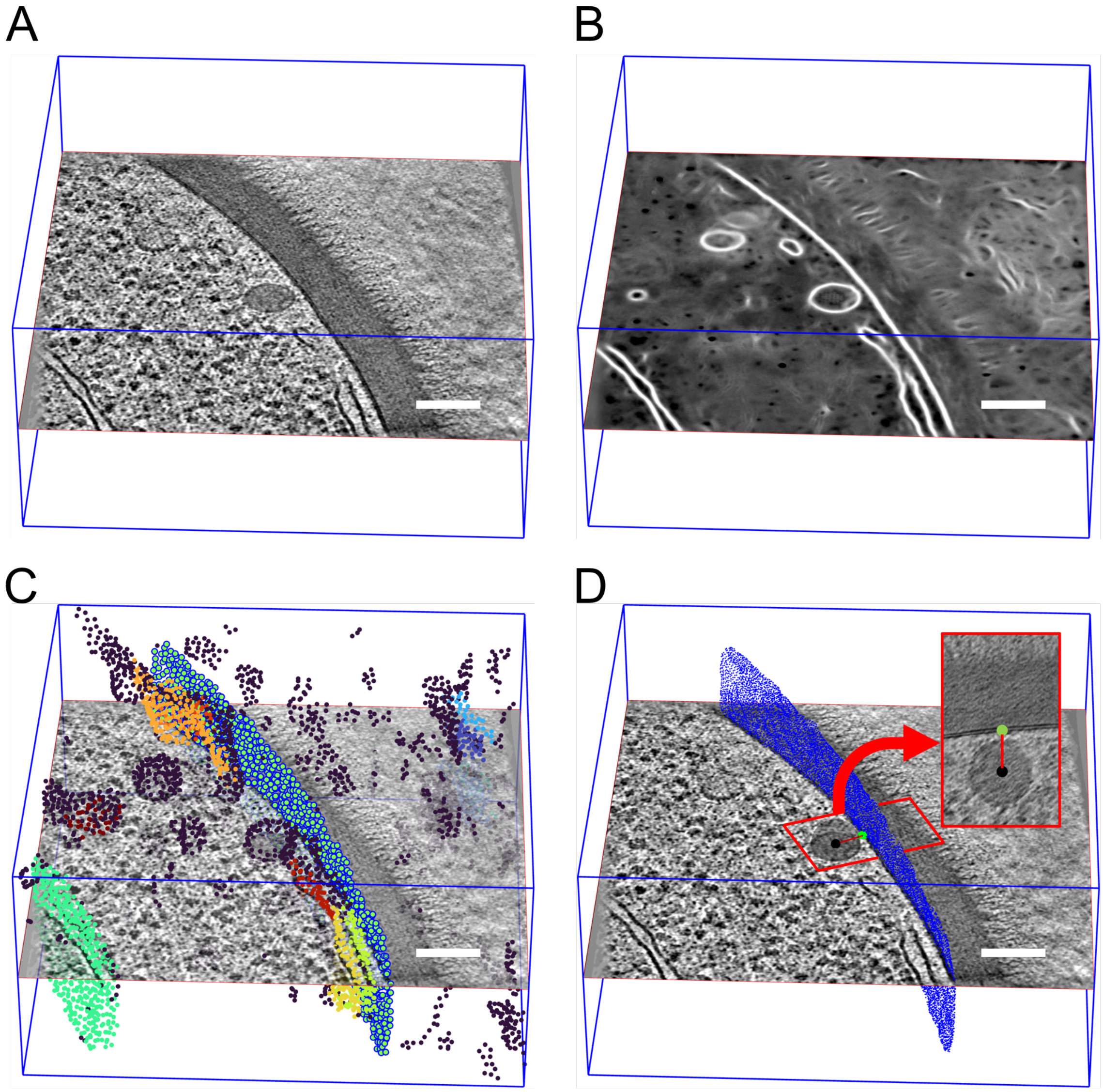


**Figure S1. Workflow of vesicle-PM distance estimation**

(A) Illustrative tomogram of an exocytic event of a yeast cell, containing a secretory vesicle.

(B) Automatic segmentation of membranes where areas identified as membranes are shown in white.

(C) Voxels classified as membrane are used to define continuous membrane models by analyzing local curvatures. Each color identifies a different membrane model. Black points indicate areas with too high or inconsistent curvature. The model corresponding to the PM is highlighted in blue (Methods).

(D) The PM model is automatically refined (blue points). The vesicle center (black point) and radius are manually identified. The closest PM point to the vesicle (green point) is used to extract a subbox for each vesicle such that it is centered on the membrane and collinear with the vesicle-membrane vector (red line). This allows for a final fair manual distance measurement. Scale bars: 100 nm.

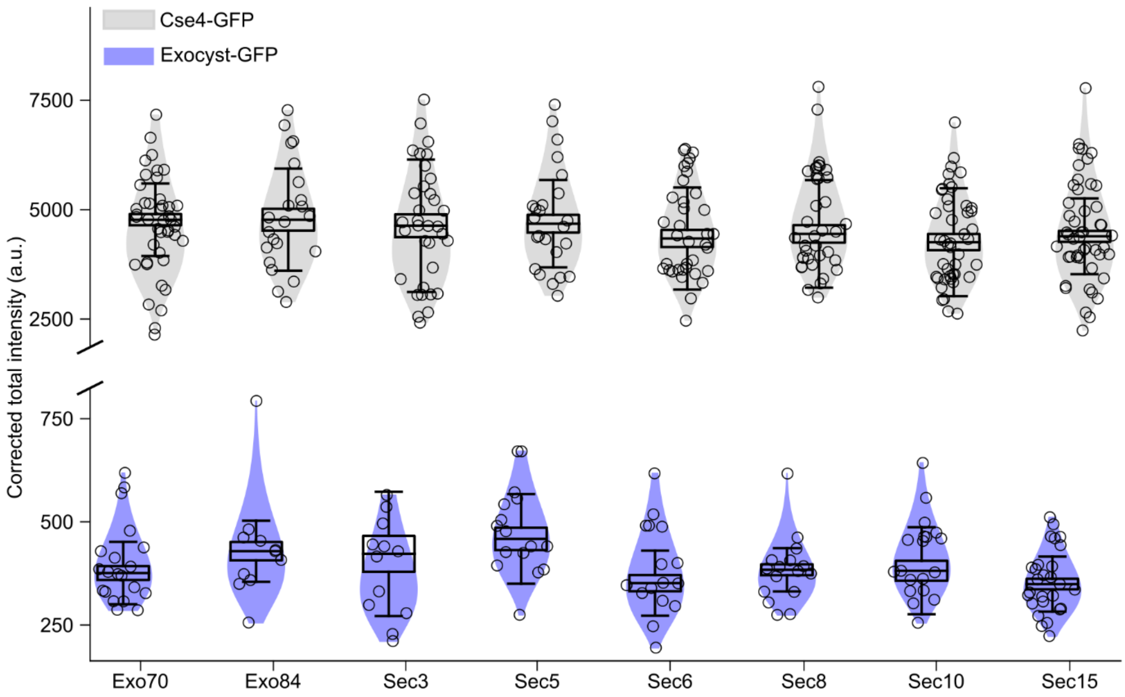


**Figure S2. Quantification of the copy number of subunits at exocyst clusters**

Cells expressing the indicated exocyst subunit fused to GFP were co-cultured with cells expressing Cse4-GFP. Upon imaging z-stacks, the copy number of each exocyst subunit fused to GFP was measured by using Cse4-GFP spots in cells in anaphase as a reference (Methods). The height of the boxes and the length of the whiskers represent the SEM and the SD (Methods). Number of exocyst subunit/Cse4 clusters analyzed (n): 20/42 (Exo70), 17/20 (Exo84), 16/34 (Sec3), 18/24 (Sec5), 21/36 (Sec6), 22/38 (Sec8), 23/44 (Sec10) and 27/48 (Sec15).

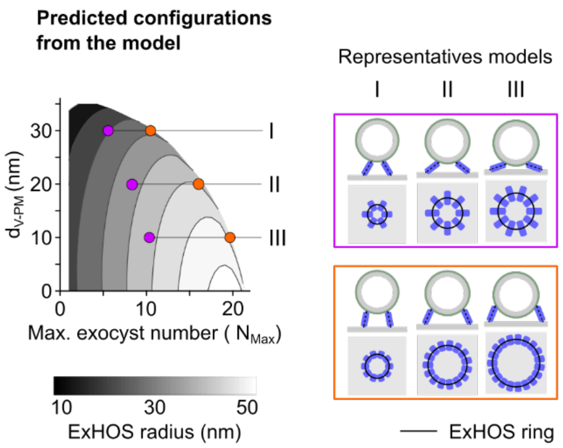


**Figure S3. Modeling of possible ring-shaped ExHOS during vesicle tethering**

The ExHOS radius is plotted according to the maximal number of exocyst that can fit in it and the distance between the PM and the vesicle surface (left). Representative solutions for the configurational space are shown (right).

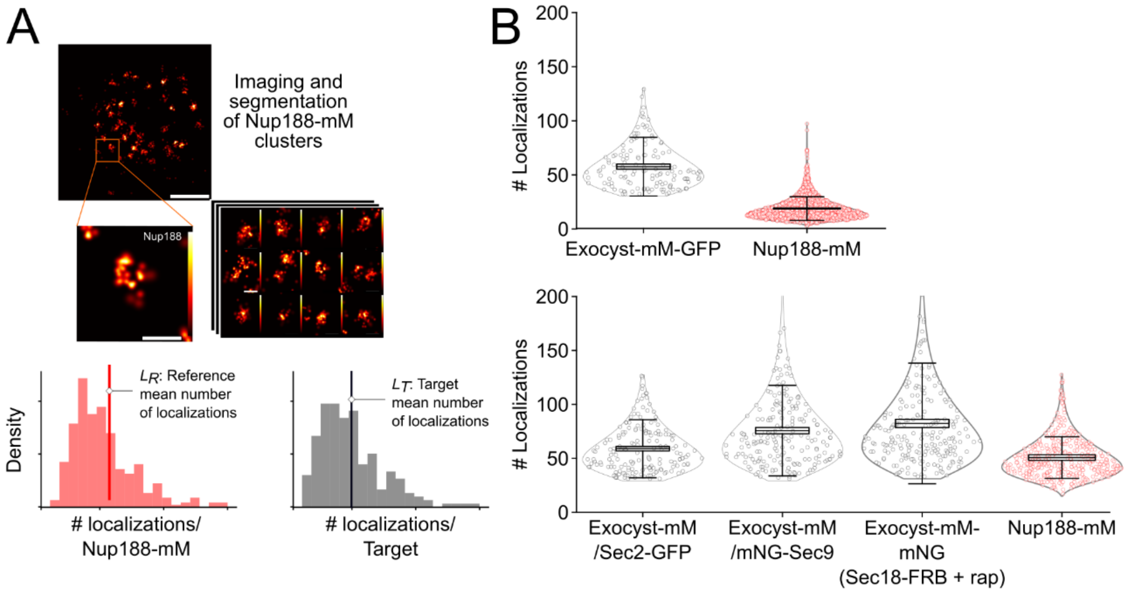


**Figure S4. Quantification of exocyst copy number in SMLM datasets**

(A) Representative images of Nup188-mM clusters, which were used as reference to determine the number of exocysts populating the clusters (Methods). The exocyst copy number, $N_T$, is calculated as $N_T = (L_T/L_R) * N_R/3$, where $L_T$ and $L_R$ are the mean number of localizations per Nup188-tmM cluster and exocyst-mM cluster, respectively. $N_R$ = 16 is the average number of Nup188 copies in the nuclear pore complex and the factor 3 is added to take into account that each exocyst is labeled with 3 mMaples. (B) A comparison of the number of localizations between the reference distribution (Nup188-mM) and exocyst-mM clusters for each sample is shown. The height of the boxes and the length of the whiskers represent the pooled SEM and the pooled SD, for three biological replicates, respectively (Methods).

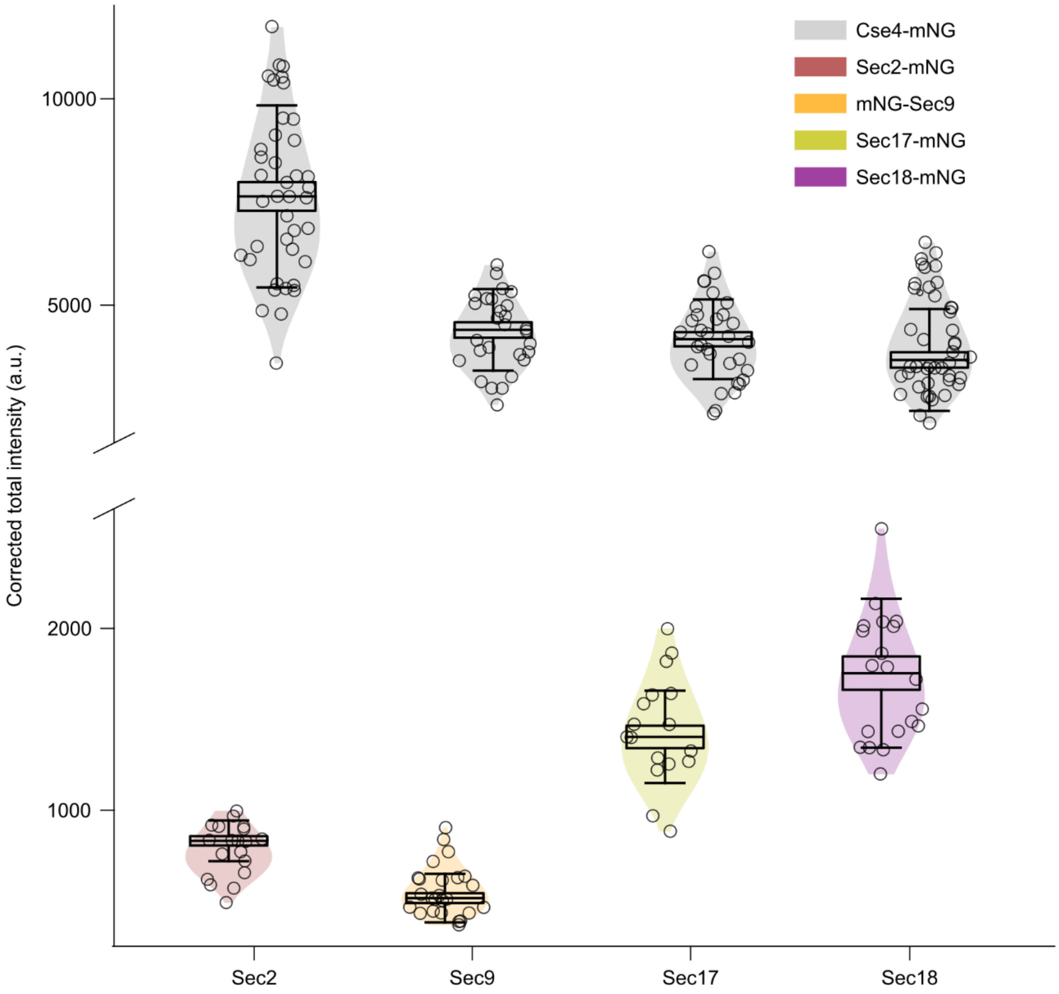


**Figure S5. Quantification of copy numbers within Sec2-mNG, mNG-Sec9, Sec17-mNG and Sec18-mNG clusters**

Cells expressing Sec2, Sec9, Sec17 or Sec18 fused to mNG were co-cultured with cells expressing Cse4-mNG. Upon imaging z-stacks, the copy number of each protein fused to mNG was measured by using Cse4-mNG spots in cells in anaphase as a reference (Methods). The height of the boxes and the length of the whiskers represent the SEM and the SD, (Methods). Number of NG-fusion/Cse4 clusters analyzed (n): 19/40 (Sec2), 25/28 (Sec9), 17/32 (Sec17) and 20/44 (Sec18).

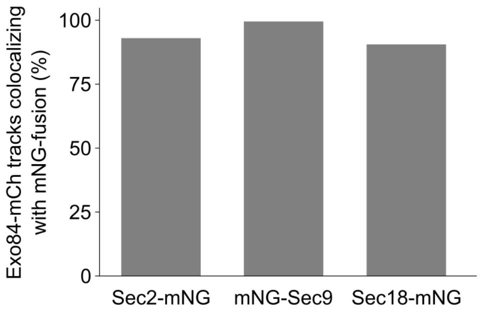


**Figure S6. Colocalization of Exo84-mCh tracks with NG-fused proteins**

For at least 90% of the tracks of Exo84-mCh clusters colocalization could be detected with Sec2-mNG (n = 31), mNG-Sec9 (n = 32) and Sec18-mNG (n = 11).

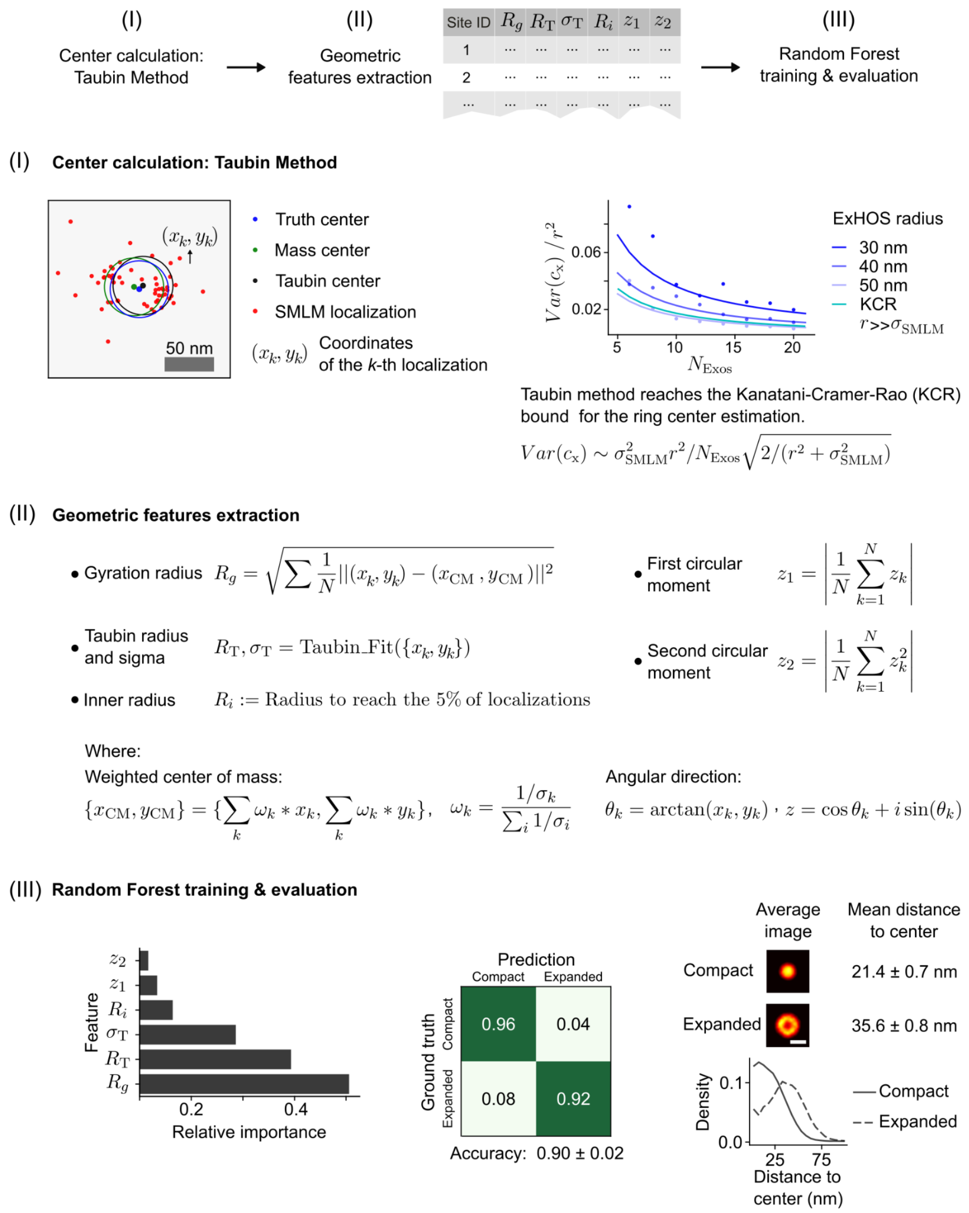


**Figure S7. Training workflow for the random forest classifier of exocyst clusters imaged by SMLM**

The pipeline used to train a random forest classifier is schematized and divided in 3 steps.

(I) The center of synthetic SMLM images corresponding to the modeled ExHOS (Methods) is calculated using the Taubin method, which reaches the minimum possible *KCR* bound. As expected, the resulting *KCR* bound depends on the density of exocysts (number of exocysts/ring radius) and is limited by the resolution of the imaging ($\sigma_{SMLM}$). The right scatter plot shows the error associated to the Taubin center estimation in synthetic SMLM images of ExHOS with varying radii and number of exocysts ($N_{Exos}$). The theoretical distribution of errors is shown with continuous lines.

(II) Synthetic SMLM images representing the modeled ExHOS are described by a set of geometrical parameters, including the gyration radius of localizations ($R_g$), the radius and its associated error ($R_T$, $\sigma_T$), the inner radius ($R_i$), and the first ($z_1$) and second ($z_2$) circular moment of the localizations. Except for the gyration radius, all the features are calculated based on the Taubin center.

(III) A random forest classifier is trained with the values of the geometrical features to distinguish SMLM images of Compact and Expanded ExHOS. The relative importance (left) of each feature and the confusion matrix (middle) are shown. Finally, the average image, the resulting radial distribution and average mean localization distance to center are shown for the resulting classification of the synthetic dataset (right). Scale bar: 100 nm.

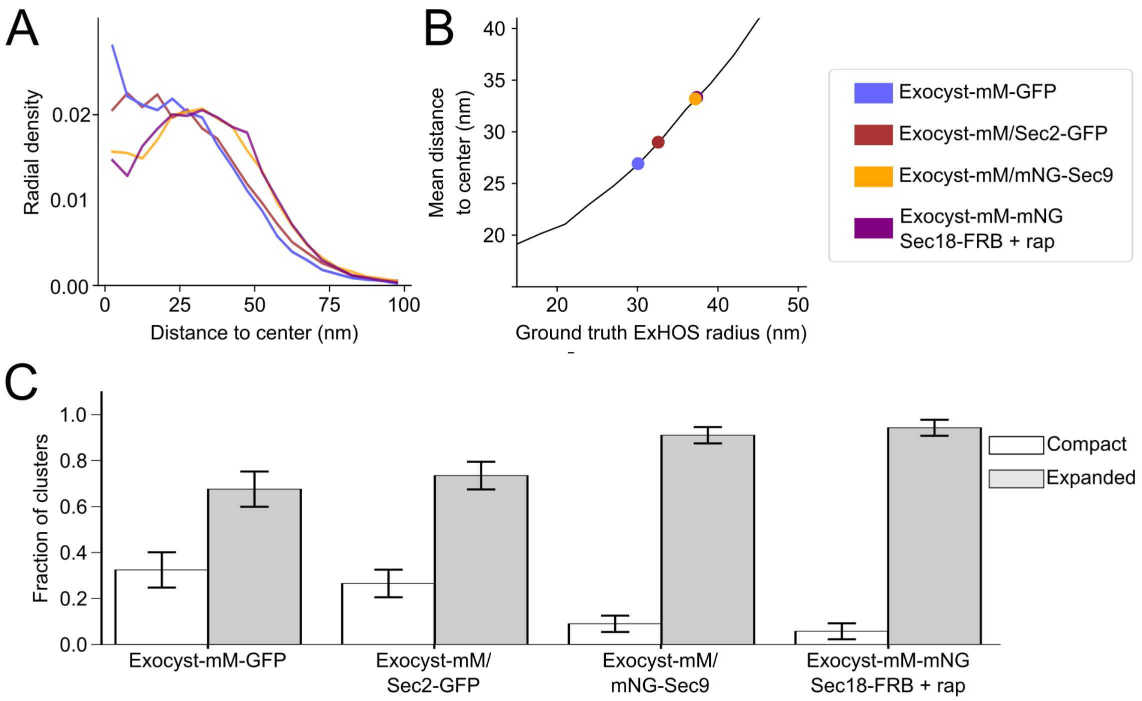


**Figure S8. Characterization of the exocyst clusters by SMLM**

(A) Radial distribution of localizations in the average images of each sample. Due to the resolution limit of SMLM and the fluorescent label size, the spatial distribution of fluorescent localizations deviates from the underlying exocyst positions in a cluster.

(B) To aid the accurate reconstructions of the ExHOS we used a synthetic dataset to generate a calibration curve (black line) describing the relationship between the measured mean distance-to-center of the localizations in SMLM images and the underlying ExHOS radius. The ExHOS radius was estimated by calculating the mean distance to the center from the radial distributions in (A) and comparing these values to the calibration curve.

(C) Fraction of the exocyst clusters belonging to the Compact and Expanded classes for each of the samples. Error bars represent the pooled SEM for three biological replicates.

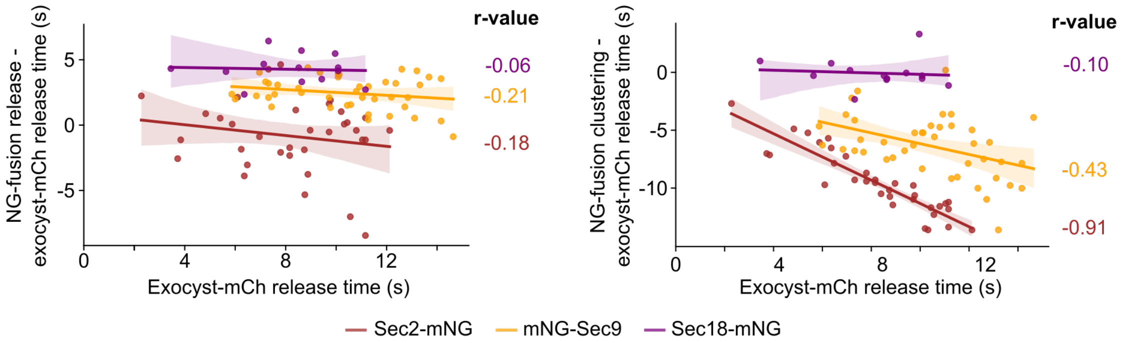


**Figure S9. Characterization of the 2-color tracking results**

Correlation between the release of exocyst-mCh and the release (left) and the clustering (right) of the corresponding mNG-fused protein. n = 34 (Sec2-mNG, brown), 42 (mNG-Sec9, orange), and 13 (Sec18-mNG, purple) tracks.

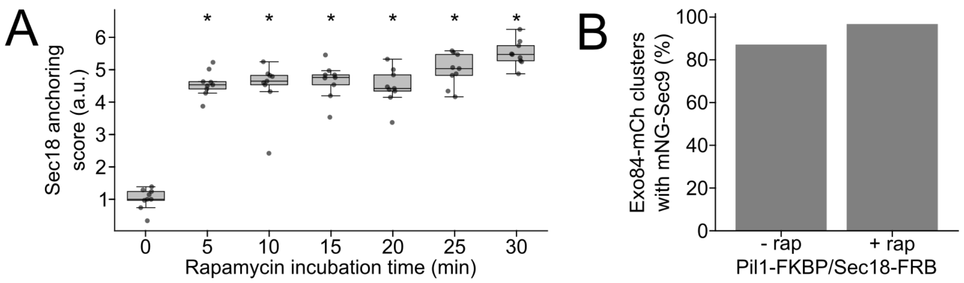


**Figure S10. Characterization of Sec18-anchor-away**

(A) Sec18-FRB-GFP recruitment to Pil1-RFP-FKBP anchors after incubating the cells with 10 µM rapamycin for the indicated times. Recruitment was assessed by quantifying the colocalization between Sec18-FRB-GFP and Pil1-RFP-FKBP foci. Recruitment scores were normalized to basal conditions (0 minutes of rapamycin incubation time). n = 9, per timepoint. * = $p < 0.05$.

(B) Exocyst-mCh clusters were assessed for the presence and absence of mNG-Sec9 clusters within their lifetimes in cells that had been treated with 10 µM of rapamycin for 10 minutes to induce the depletion of Sec18-FRB from exocytic events (n = 20). No defect in the recruitment of mNG-Sec9 to exocyst-mCh clusters could be detected upon Sec18-FRB anchoring-away.

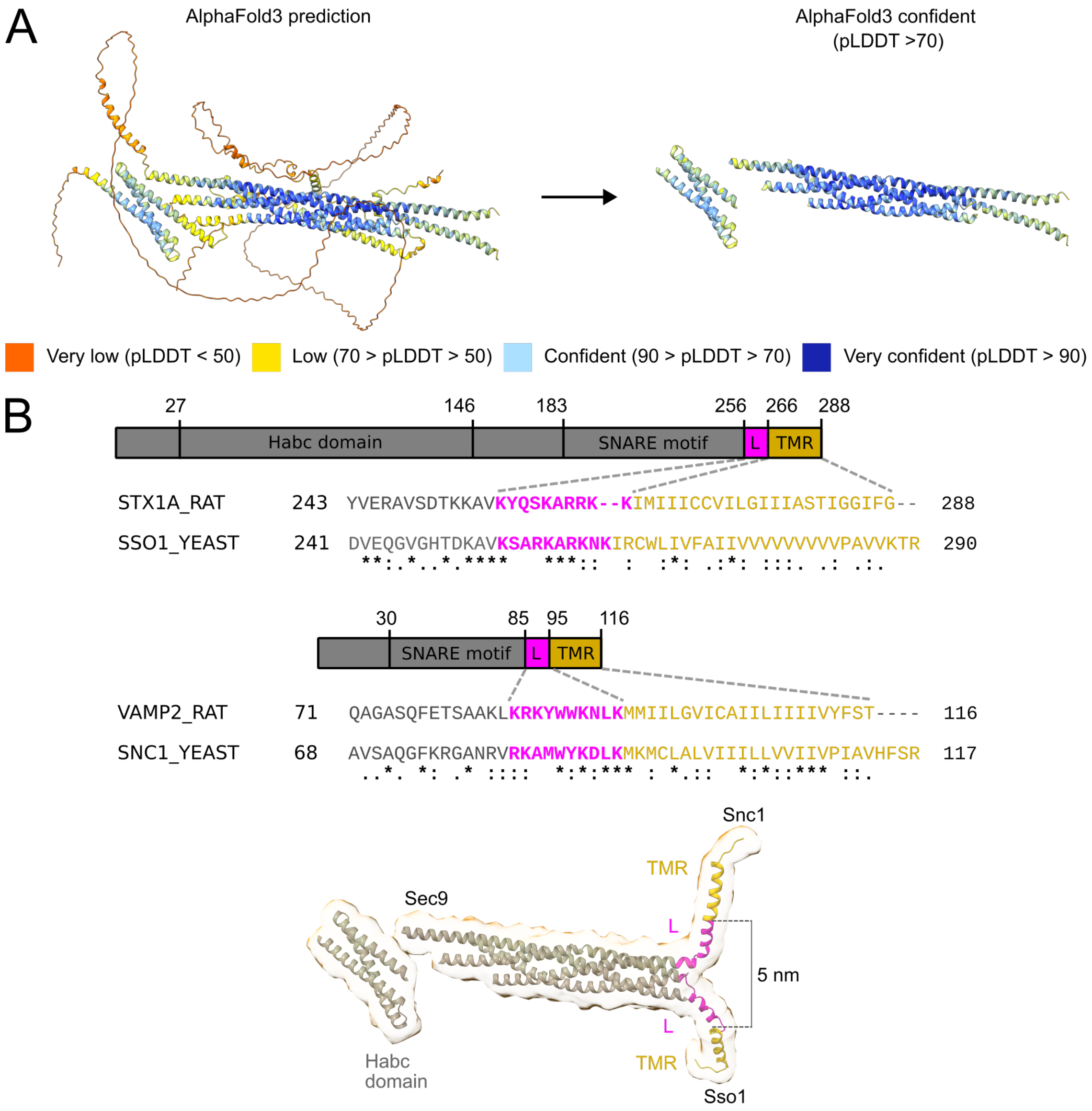


**Figure S11. Modeling the yeast exocytic *trans*-SNARE complex**

(A), The structure of the *S. cerevisiae cis*-SNARE complex was predicted with AlphaFold3 (left). AlphaFold3 prediction regions with pLDDT > 70 were used to represent the confident ribbon diagram of the *cis*-SNARE complex (right).

(B), Sequence alignment between the *R. novergicus* SNARE proteins, Syntaxin1 (STX1A_RAT) and Vamp2 (VAMP2_RAT), and their respective *S. cerevisiae* homologs, Sso1 (SSO1_YEAST) and Snc1 (SNC1_YEAST) in *S. cerevisiae* sequences (top and middle). Illustration of the modeled *trans*-SNARE complex (bottom). The short juxtamembrane linker (L, magenta) and the C-terminal transmembrane region (TMR, gold) are highlighted.

| Dataset | Vesicle ID | $d_{V\text{-}PM}$ (nm) |
|---|---|---|
| Exocyst-mNG | 1 | 2.0 |
| Exocyst-mNG | 2 | 3.6 |
| Exocyst-mNG | 3 | 4.3 |
| Exocyst-mNG | 4 | 4.5 |
| Exocyst-mNG | 5 | 4.6 |
| Exocyst-mNG | 6 | 6.4 |
| Exocyst-mNG | 7 | 7.7 |
| Exocyst-mNG | 8 | 9.9 |
| Exocyst-mNG | 9 | 11.1 |
| Exocyst-mNG | 10 | 16.4 |
| Exocyst-mNG | 11 | 17.6 |
| Exocyst-mNG | 12 | 19.8 |
| Exocyst-mNG | 13 | 27.3 |
| Exocyst-mNG | 14 | 27.4 |
| Exocyst-mNG | 15 | 37.3 |
| mSc3-Sec9 | 1 | 4.0 |
| mSc3-Sec9 | 2 | 4.1 |
| mSc3-Sec9 | 3 | 4.6 |
| mSc3-Sec9 | 4 | 5.4 |
| mSc3-Sec9 | 5 | 6.3 |

**Table S1. Vesicle to PM distances measured in cryo-CLEM tomograms**

| Strain | Genotype | Source |
|---|---|---|
| OGY0997 | *MATa, his3Δ1, leu2Δ0, met15Δ0, ura3Δ0::ADH1pr-OsTIR1-9xMyc-URA3,SEC6-mMaple::hphNT1, SEC8-mMaple::hphNT1, Sec5-mMaple::LEU2, SEC2-3xmyeGFP::kanMX4* | This study |
| OGY1045 | *MATa, his3Δ1, leu2Δ0, met15Δ0, ura3Δ0, GAL1pr-I-SCE1::HIS3, mNeonGreen-SEC9* | This study |
| OGY1056 | *MATa, his3Δ1, leu2Δ0, met15Δ0, ura3Δ0, SEC6-mMaple::hphNT1, SEC8-mMaple::hphNT1, SEC5-mMaple::KlLEU2, EXO84-3xmyeGFP::kanMX4* | This study |
| OGY1074 | *MATa, his3Δ1, leu2Δ0, met15Δ0, ura3Δ0, tor1-1, SEC2-mNeonGreen::kanMX4* | This study |
| OGY1089 | *MATa, his3Δ1, leu2Δ0, met15Δ0, ura3Δ0, PIL1-RFP-FKBP::natNT2, SEC18-FRB-GFP::kanMX4* | This study |
| OGY1093 | *MATa, his3Δ1, leu2Δ0, met15Δ0, ura3Δ0, tor1-1, SEC18-mNeonGreen::kanMX4* | This study |
| OGY1121 | *MATa, his3Δ1, leu2Δ0, met15Δ0, ura3Δ0, tor1-1, SEC17-mNeonGreen::kanMX4* | This study |
| OGY1154 | *MATa, his3Δ1, leu2Δ0, ura3Δ0, can1Δ::STE2pr-LEU2, lyp1Δ::, EXO84-3xmCherry::natNT2, SEC2-mNeonGreen::kanMX4* | This study |
| OGY1158 | *MATa, his3Δ1, leu2Δ0, ura3Δ0, can1Δ::STE2pr-LEU2, lyp1Δ::, EXO84-3xmCherry::natNT2, SEC18-mNeonGreen::kanMX4* | This study |
| OGY1200 | *MATa, his3Δ1, leu2Δ0, met15Δ0, ura3Δ0, tor1-1, can1Δ::STE2pr-LEU2, lyp1Δ::, GAL1pr-I-SCE1::HIS3, mNeonGreen-SEC9, EXO84-3xmCherry::natNT2* | This study |
| OGY1240 | *MATa, his3Δ1, leu2Δ0, met15Δ0, ura3Δ0, SEC6-mMaple::hphNT1, SEC8-mMaple::hphNT1, Sec5-mMaple::KlLEU2, tor1-1, fpr1Δ::, PIL1-FKBP::natNT2, SEC18-FRB:KlURA, EXO84-mNeonGreen::kanMX4 Kan* | This study |
| OGY1269 | *MATa, his3Δ1, leu2Δ0, ura3Δ0, LYS+, can1Δ::STE2pr-LEU2, lyp1Δ::, tor1-1, fpr1Δ::, PIL1-FKBP::natNT2, SEC18-FRB::hphNT1, GAL1pr-I-SCE1::HIS3, mNeonGreen-SEC9, EXO84-3xmCherry::KlURA3* | This study |
| OGY1320 | *MATa, his3Δ1, leu2Δ0, met15Δ0, ura3Δ0, SEC6-mMaple::hphNT1, SEC8-mMaple::hphNT1, SEC5-mMaple::KlLEU2, tor1-1, fpr1Δ::, PIL1-FKBP::natNT2, GAL1pr-I-SCE1::HIS3, mNeonGreen-SEC9* | This study |
| OGY1348 | *MATa, his3Δ1, leu2Δ0, met15Δ0, ura3Δ0, tor1-1, EXO84-2xmNeonGreen::kanMX4, SEC5-2xmNeonGreen::kanMX4, EXO70-2xmNeonGreen::hphNT1* | This study |
| OGY1490 | *MATa, his3Δ1, leu2Δ0::GAL1pr-I-SCE1::HIS3, met15Δ0, ura3Δ0, 3xmScarlet3-SEC9* | This study |
| OGY1495 | *MATa, his3Δ1, leu2Δ0, lys2Δ0, ura3Δ0, Cse4-mNeonGreen* | This study |
| OGY1539 | *MATa, his3Δ1, leu2Δ0, met15Δ0, ura3Δ0, SEC3-GFP::HIS3MX6* | GFP collection |
| OGY1540 | *MATa, his3Δ1, leu2Δ0, met15Δ0, ura3Δ0, SEC5-GFP::HIS3MX6* | GFP collection |
| OGY1541 | *MATa, his3Δ1, leu2Δ0, met15Δ0, ura3Δ0, SEC6-GFP::HIS3MX6* | GFP collection |
| OGY1542 | *MATa, his3Δ1, leu2Δ0, met15Δ0, ura3Δ0, SEC8-GFP::HIS3MX6* | GFP collection |
| OGY1543 | *MATa, his3Δ1, leu2Δ0, met15Δ0, ura3Δ0, SEC10-GFP::HIS3MX6* | GFP collection |
| OGY1544 | *MATa, his3Δ1, leu2Δ0, met15Δ0, ura3Δ0, SEC15-GFP::HIS3MX6* | GFP collection |
| OGY1545 | *MATa, his3Δ1, leu2Δ0, met15Δ0, ura3Δ0, EXO70-GFP::HIS3MX6* | GFP collection |
| OGY1546 | *MATa, his3Δ1, leu2Δ0, met15Δ0, ura3Δ0, EXO84-GFP::HIS3MX6* | GFP collection |
| OGY1547 | *MATa, his3Δ1, leu2Δ0, met15Δ0, ura3Δ0, CSE4-GFP::HIS3MX6* | GFP collection |
| yPH0008 | *MATa, his3 200, leu2-3,112, ura3-52, lys2-801, NUP188-mMaple::HIS3MX6* | Ries' lab |

**Table S2. List of strains used in this study**
